# Chrysin directing an enhanced solubility through the formation of a supramolecular cyclodextrin-calixarene drug delivery system: a potential strategy in antifibrotic diabetes therapeutics

**DOI:** 10.1101/2023.10.03.560552

**Authors:** Anca Hermenean, Eleftheria Dossi, Alex Hamilton, Maria Consiglia Trotta, Marina Russo, Caterina Claudia Lepre, Csilla Sajtos, Ágnes Rusznyák, Judit Váradi, Ildikó Bácskay, István Budai, Michele D’Amico, Ferenc Fenyvesi

## Abstract

Calixarene 0118 (OTX008) and chrysin (CHR) are promising molecules for the treatment of fibrosis and diabetes complications but require an effective delivery system to overcome their low solubility and bioavailability. Sulfobutylated β-cyclodextrin (SBECD) was evaluated for its ability to increase the solubility of CHR by forming a ternary complex with OTX008. The resulting increase in solubility and the mechanisms of complex formation were identified through phase-solubility studies, while dynamic light-scattering assessed the molecular associations within the CHR-OTX008-SBECD system. Nuclear magnetic resonance, differential scanning calorimetry, and computational studies elucidated the interactions at the molecular level, and cellular assays confirmed the system’s biocompatibility. Combining SBECD with OTX008 enhances CHR solubility more than using SBECD alone, by forming water-soluble molecular associates in a ternary complex. This aids in the solubilization and delivery of CHR and OTX008. Structural investigations revealed non-covalent interactions essential to complex formation, which showed no cytotoxicity in hyperglycemic *in vitro* conditions. A new ternary complex has been formulated to deliver promising antifibrotic agents for diabetic complications, featuring OTX008 as a key structural and pharmacological component.

## 1. Introduction

Calixarenes and cyclodextrins are cyclic molecules, with synthetic and natural origin respectively and they differ in their building blocks [1]. Calixarenes are synthetic cyclic molecules built from phenolic units linked by methylene bridges, which can be made water-soluble with hydrophilic modifications [2], whereas cyclodextrins are naturally occurring water-soluble cyclic oligosaccharides composed of glycosidic units connected by α-1,4-glucosidic bonds [3].

Both calixarenes and cyclodextrins share the characteristic of being able to encapsulate guest molecules within their structures, which can enhance the solubility and bioavailability of drugs that are otherwise poorly soluble [2,3]. Specifically, water-soluble calixarenes, like 4-sulphonic calix[n]arenes, have the capacity to increase the solubility of testosterone in water, with effectiveness dependent on their ring size [4]. Previous studies have demonstrated that β-cyclodextrins, including sulfobutylated β-cyclodextrin (SBECD), are effective at improving the solubility and permeability of chrysin (CHR) [5]. CHR is a bioflavonoid, present in the honey and propolis and has a limited bioavailability, due to its poor water-solubility [6]. Recognized for its various therapeutic properties, CHR exhibits anti-inflammatory, anticancer, and antioxidant activities [7,8].

Recent studies have also highlighted its potential antifibrotic effects [9–11]. OTX008 has the capability to bind with biomolecules like proteins and nucleic acids, influencing enzyme functions [12]. It exhibits anti-cancer properties by suppressing cancer cell growth and tumor angiogenesis [13]. Zucchetti et al. observed that OTX008, a galectin-1 (Gal-1) targeting compound, can inhibit endothelial cell activities including proliferation and motility [14]. Additionally, OTX008 has been found to inhibit the overproduction of Gal-1, a protein abundant in the kidneys of diabetic mice and a significant contributor to fibrosis in diabetes [15]. By inhibiting Gal-1 accumulation, OTX008 showcases as a new therapeutic inhibitor of Gal-1, aiming to address fibrosis in diabetes.

Even if both OTX008 and CHR are promising molecules for the treatment of fibrosis acting on different targets they have limited absorption and poor water solubility and, thus a suitable drug delivery system (DDS) is required for their combined therapeutic application. The common formulation and delivery of the water insoluble flavonoid CHR and calixarene OTX008 has not been reported yet, therefore the aim of the research was to investigate their joint delivery by the following strategy.

While the molecular interaction between cyclodextrins and CHR is well-documented [5], a similar relationship of CHR with OTX008 has not been reported yet. The combined use of calixarenes and cyclodextrins to enhance solubility remains largely unexplored. The water solubility of niclosamide was previously enhanced through the joint use of 4-sulphonato-calix[6]arene and hydroxypropyl-β-cyclodextrin [16]. Considering OTX008’s small cavity and limited water solubility, a weak interaction with CHR was anticipated.

Our strategy entailed employing SBECD to improve the solubility of OTX008, enhancing its drug delivery capabilities.

We aimed at assessing the ability of these macrocycles, both individually and in combination, to complex with CHR, forming an advanced DDS. This involved analyzing the phase-solubility of CHR with SBECD in binary arrangements, as well as the phase-solubility in ternary systems involving CHR with OTX008+SBECD. The molecular structures of these compounds are presented in Figure 1.

**Figure 1.**
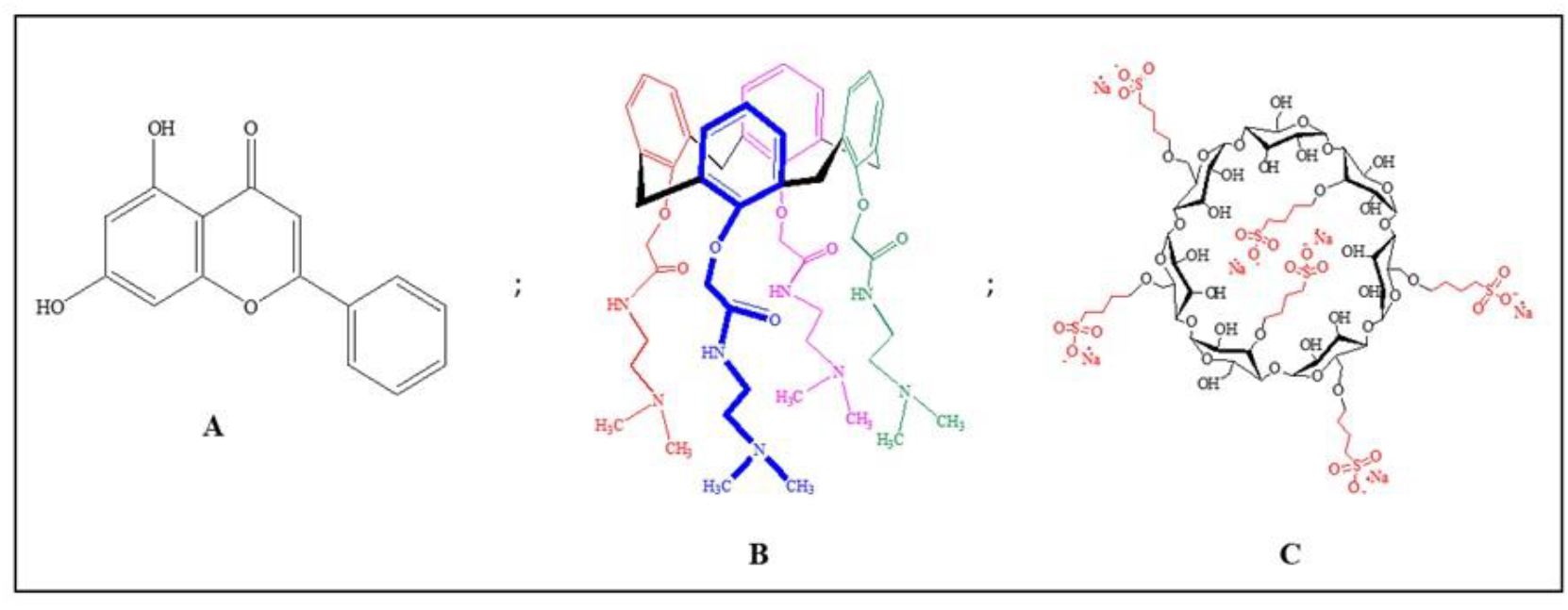
Chemical structures of (A) CHR, (B) OTX008 and (C) SBECD.

The selection of SBECD was based on its anionic nature. Given their structural attributes, we anticipated the formation of supramolecular complexes through the interactions of OTX008 and anionic SBECD with CHR, mediated by non-covalent bonds. To examine this hypothesis, we measured the size distribution of the water-based complexes using dynamic light scattering (DLS). We conducted chemical (NMR) and thermal (DSC) analyses of the individual components and their binary mixtures to understand the interactions within the ternary mix, complemented by molecular modelling.

The synthesized complexes were also subjected to *in vitro* biocompatibility assays using embryonic rat cardiac H9c2 cells in both normal and high-glucose conditions, which established a foundation for their potential use in medical treatments.

## 2. Results

### 2.1. OTX008 solubilization with SBECD

Due to its low water solubility, OTX008 required the use of SBECD (Figure 1.) as solubilizing agents to reach the therapeutically relevant concentration of 0.75 mg/ml. SBECD proved to be an effective solubilizer; solutions of 7.3 mass/mass concentration (m/m%) SBECD was successful in dissolving OTX008 to the target concentration in water. These solution concentrations were therefore utilized for subsequent experiments.

### 2.2. CHR solubilization in OTX008-SBECD solution

While CHR (Figure 1.) does not dissolve in water, its solubility can be enhanced by cyclodextrins, as previously reported [5]. In this study, we examined the solubility of CHR in the solution of 7.3 m/m% SBECD with and without the addition of 0.75 mg/ml OTX008. We observed a significant increase in CHR solubility in the presence of OTX008 (p<0.0001) when compared to solutions containing only cyclodextrins, as shown in Figure 2.

**Figure 2.**
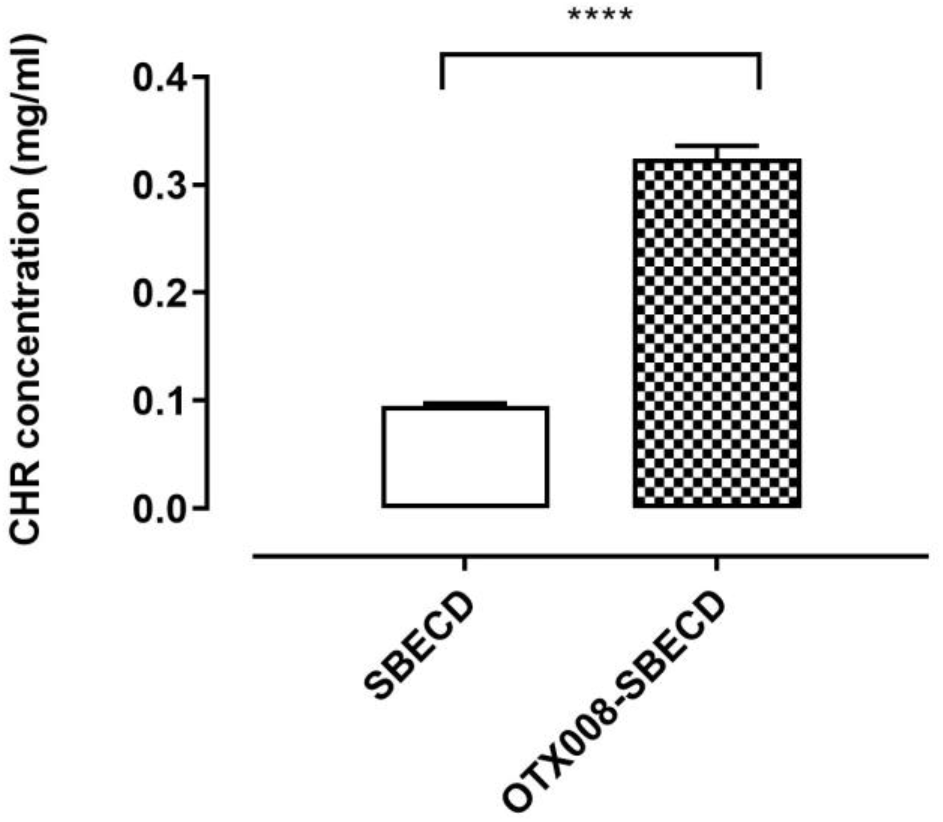
CHR is solubilized in water by SBECD (7.3 m/m%), and OTX008-SBECD (OTX008 concentration: 0.75 mg/ml). The simultaneous application of OTX008 and SBECD significantly increased the solubility of CHR compared to SBECD (n=3; ****p<0.0001).

After clarification, the solutions were lyophilized to yield solid products, which were then reconstituted in 0.9 m/m% NaCl solution. The lyophilized products dissolved completely in the saline solution, making them suitable for further *in vitro* tests and *in vivo* experimentation with parenteral (intravenous or intraperitoneal) administration. The structures of the lyophilized products are presented on Figure 3.

**Figure 3.**
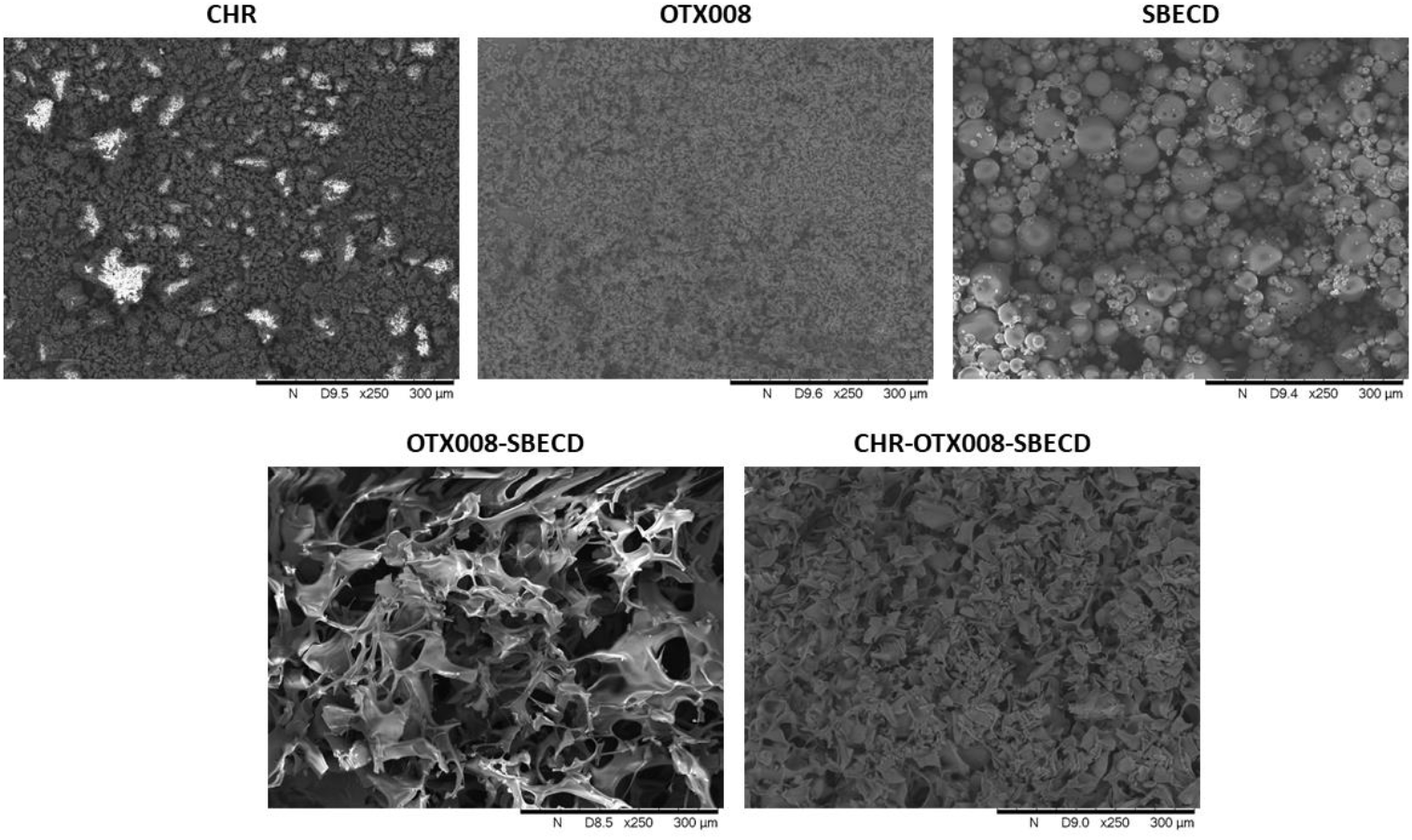
Scanning electron microscopic images of CHR, OTX008, SBECD, OTX008-SBECD and CHR-OTX008-SBECD. Both OTX008-SBECD and CHR-OTX008-SBECD have a porous structure after lyophilization and differs from the pure raw materials. (The white regions show the accumulation of charges on projecting structures.)

### 2.3. Phase-solubility study

During the phase-solubility analysis, a range of dilutions for SBECD and OTX008-SBECD, solutions were employed to dissolve CHR. The solubility of CHR corresponded linearly with the concentrations of SBECD. In contrast, the OTX008-SBECD solutions exhibited non-linear, positively skewed curves, indicating the formation of associations in the solutions (Figure 4.).

**Figure 4.**
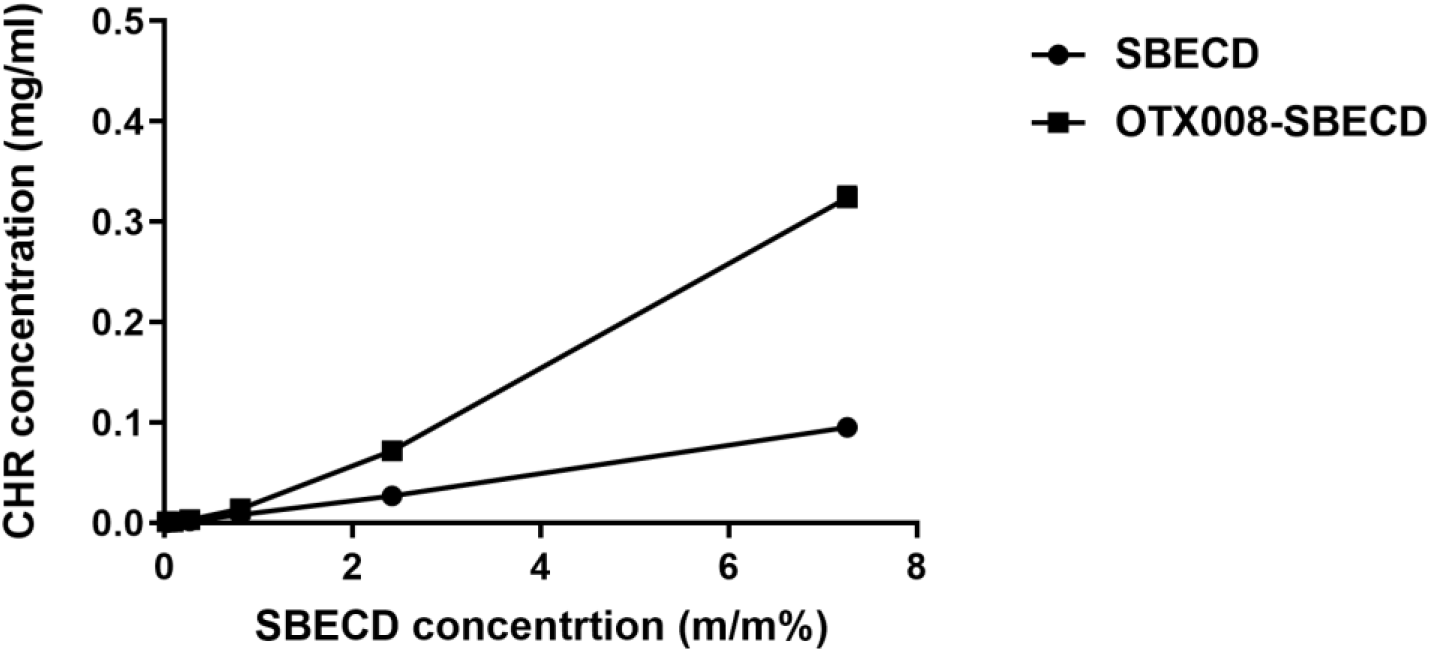
Phase-solubility curves of CHR in SBECD and OTX008-SBECD solutions. OTX008 concentration was 0.75 mg/ml in the solutions with the highest SBECD concentrations at 7.3 m/m% SBECD.

### 2.4. Size distribution measurement of the cyclodextrin complexes with dynamic light scattering (DLS)

When measured in solution at a concentration of 7.3 m/m%, SBECD displayed molecule associations with a particle size of 351.8 nm as per Dynamic Light Scattering (DLS) data. The particle size was found to increase upon mixing SBECD with OTX008, registering at 1002 nm, and it further escalated significantly higher, than 2000 nm (and even higher, which size range is out of the detection of the instrument) upon the addition of CHR, forming the ternary CHR-OTX008-SBECD complex (Table 1 and Figure S1).

**Table 1.** Size distribution of SBECD, CHR-SBECD, OTX008-SBECD, and CHR-OTX008-SBECD complexes.

|  | Peak 1. |  | Peak 2. |  | Peak 3. |  |
| --- | --- | --- | --- | --- | --- | --- |
|  | Size | Intensity % | Size | Intensity % | Size | Intensity % |
| SBECD | 351.8 nm | 100 % | - | - | - | - |
| CHR-SBECD | 1.1 nm | 20.8 % | 94.83 nm | 38.1% | 3764 nm | 41.1% |
| OTX008-SBECD | 1.5 nm | 13.2% | 1002 nm | 86.8% | - | - |
| CHR-OTX008-SBECD | 1.2 nm | 24.8% | >2000 nm | >70% | - | - |

### 2.5. pH-dependent CHR solubility determination

The influence of pH on the solubility of CHR, with and without the presence of OTX008, was examined. An increase in pH led to a slightly increase in CHR solubility when 7.3 m/m% SBECD was present, as shown in Figure 5.

**Figure 5.**
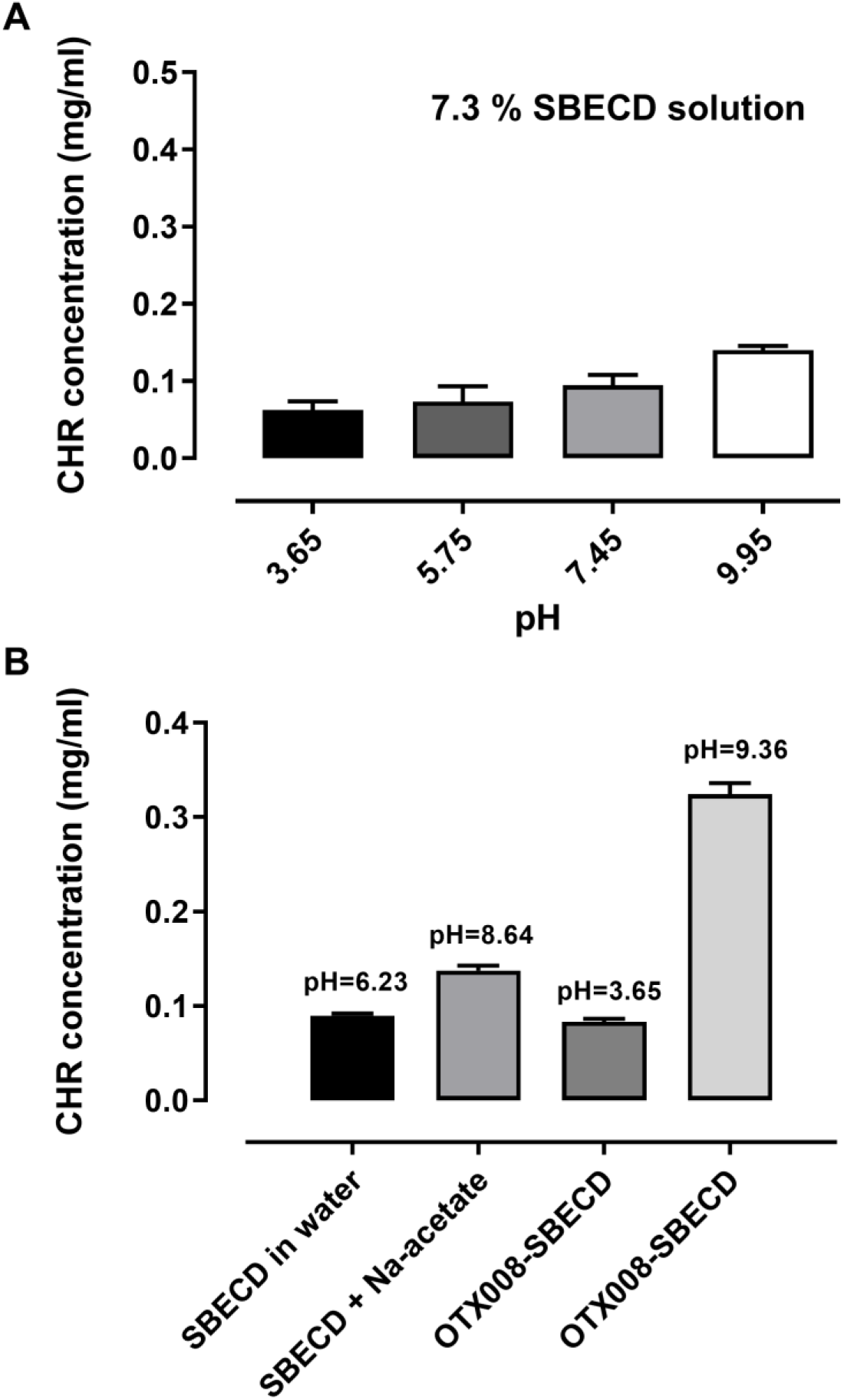
pH-dependent solubility of CHR in SBECD solution (A) and in the presence of OTX008 (B). OTX008 increased the pH of the solutions and caused increased solubility of CHR.

The initial pH of the 7.3 m/m% SBECD solution was 6.23. Adjusting the pH of the SBECD solution to 8.64 using sodium acetate resulted in a slight improvement in CHR solubility. Conversely, acidifying the OTX008-SBECD solution caused a notable reduction in CHR solubility, aligning it with the levels observed in the SBECD solution alone. Nevertheless, when OTX008 was combined with SBECD in alkaline conditions, CHR solubility was significantly enhanced, increasing three to fourfold. The pH level of the OTX008-SBECD solution was 9.36, when prepared in purified water.

Changing the pH from 3.65 to 9.95 caused only a twofold improvement of CHR’s solubility in the presence of SBECD. Both OTX008 and alkalinity are necessary for the significant solubility improvement with SBECD. In the presence of OTX008 and SBECD the pH changes to 9.36 resulted a fourfold increase in the solubility of CHR.

Ternary complexes are formed between CHR, OTX008, and SBECD, leading to an increase in the size of molecular associations. The interaction of OTX008 with SBECD and the subsequent formation of a ternary complex are essential for the notable increase in CHR solubility.

### 2.6. NMR studies

To elucidate the reasons behind the enhanced solubility of CHR in presence of both OTX008 and SBECD, ^1^H and Nuclear Overhauser Effect Spectroscopy (NOESY) NMR spectra of the ternary mixture CHR-OTX008-SBECD, binary mixtures CHR-SBECD and OTX008-SBECD and single components CHR [17], OTX008 and SBECD were recorded. The ideal deuterated solvent for the spectrometric analysis of all components was DMSO-d_6_ while D_2_O used as best representative of physiological conditions and the evaluation of potential interactions between the components. The spectra of clear solutions of CHR (Figure 6A.) and OTX008 (Figure S2) were recorded in DMSO-d_6_ as from their insolubility in D_2_O. The SBECD was examined in both deuterated solvents and when it was in mixtures, showed the strongest and predominant absorption to the applied magnetic field (Figure S3 and Figure S4), due to its high concentration in the solutions.

**Figure 6.**
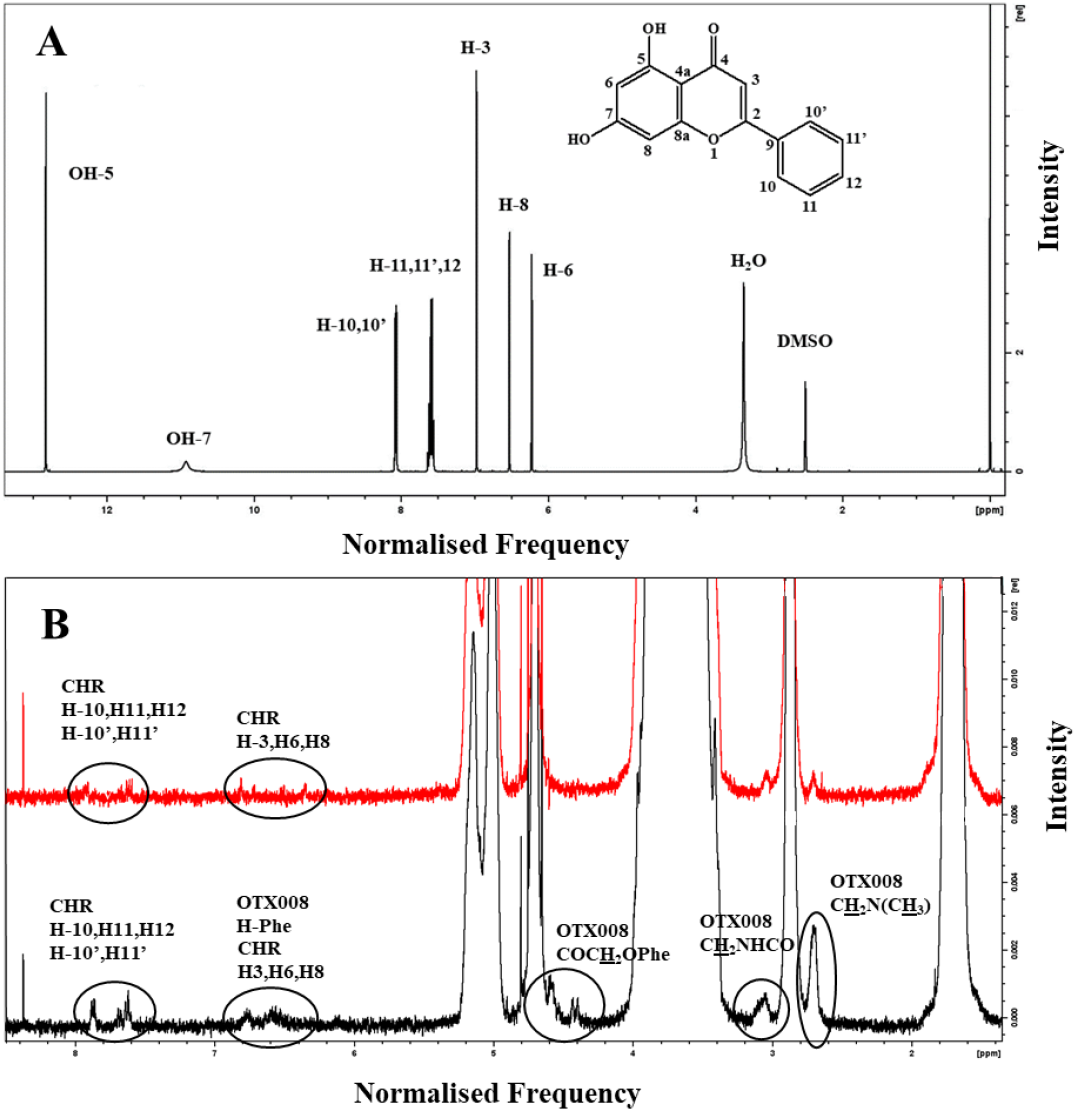
(A) Full ^1^H NMR spectrum of CHR in DMSO-d_6_ with assigned peaks. (B) ^1^H NMR spectra of CHR-OTX008-SBECD mixture (Black line) and CHR-SBECD mixture (Red line) in D_2_O showing the presence of CHR and OTX008 (Figure 1) in the ternary structure (framed areas), area 9.0-1.8 ppm. All CHR and OTX008 assignments are extrapolated from their spectra in DMSO-d_6_. All the unsigned large peaks are associated to SBECD proton absorptions (Figure 1).

#### 2.6.1. Binary mixture of CHR-SBECD

Mixing CHR with SBECD at a molar ratio of 0.01 (CHR/SBECD) in DMSO-d_6_, as seen in Figure S5, or in D_2_O, shown in Figure S6, results in ^1^H NMR spectra that are indistinguishable from that of pure SBECD. This supports the high propensity of CHR to be hosted within the toroidal cavity of the cyclodextrin structure. Such inclusion is consistent with the formation of sizable aggregates detected in DLS studies and is corroborated by the absence of free CHR signals in the NOESY spectra of the binary mixture as depicted in Figures S7 and Figure S7a.

#### 2.6.2. Binary mixture of OTX008-SBEC

The NMR spectra of the binary mixture, composed at a molar ratio of 0.02 (OTX008/SBECD), display signals corresponding only to SBECD when analyzed in DMSO-d_6_, as shown in Figures S8 and Figures S9. This observation is likely due to the chosen molar proportions of the components in the mixture. Interestingly, computational simulations, as outlined in the subsequent paragraph, indicate that the interaction between SBECD and OTX008 is not energetically favored (+0.9 kcal mol^-1^).

#### 2.6.3. Ternary mixture CHR-OTX008-SBECD

When CHR is mixed with SBECD (0.03 mol/mol, CHR/SBECD) and OTX008 (1.6 mol/mol, CHR/OTX008) in DMSO-d_6_ (Figure S10, Figure S10a - S10c and Figure S11) or D_2_O (Figure S12), the ^1^H NMR spectra of the mixture show some differences in protons chemical shifts when compared to those observed for the single components of the mixture. In DMSO-d_6_ spectra, the signals attributed to the two OHs of the CHR molecule bonded to carbon atoms in 5 and 7 position are seen very weak or even not seen at all (Figure 6A.). This can be due to the molar ratio of the CHR in the ternary mixture or because are involved in the non-bonded interaction that keeps the supramolecular structure together.

No changes observed for protons bonded to carbon atoms in positions 10 to 10’ suggesting that the potential interaction of the CHR with the other two components of the ternary mixture is involving the opposite part of the molecule; small chemical shifts to higher magnetic field, 0.02 ppm, 0.05 ppm and 0.03 ppm respectively for protons H-3, H-8 and H-6 indicating interaction of the condensed part of the CHR (Figure 6A, Figure S10b and S10c). The shape of the last 3 protons is broader that could imply electrostatic interactions through hydrogen bonding with the two other components (OTX008 and SBECD) partners which contain several of oxygen and/or nitrogen atoms.

In the same spectrum (Figure S10), OTX008 protons bonded to nitrogen atoms (NH) are subjected to a small chemical shift to a higher magnetic field from 8.32 to 8.29 ppm (0.03 ppm). Interestingly, a larger down field effect from 6.65 ppm to 6.71-6.59 ppm of OTX008 aromatic protons indicated that the upper aromatic cup shape of OTX008 is distorted by the 3 members interaction (Figure S10c). This is also supported by the broadness of the resulted signal centered at 6.70 ppm. The above observations indicated that the CHR molecules are electrostatically bonded to carbonyl groups of the OTX008 side arms and are causing this distortion (confirmed by simulations). No changes on the chemical shifts of the SBECD protons (Figure S10).

Three signals attributed to CHR aromatic protons H-3, H-8 and H-6 at 6.60 ppm, 6.55 ppm and 6.22 ppm respectively are appeared in the NOESY spectra of the ternary mixture confirming that any interaction of CHR with the system is involving the phenolic aromatic ring (Figure S11 and Figure S12).

Some small differences (0.05 ppm) in the chemical shifts of the aromatic protons of CHR were observed in spectra recorded in D_2_O when the binary CHR-SBECD and the ternary mixture were compared (Figure 6.). Those peaks were broader indicating the complex new situation in the macrostructure. The NOESY spectra also confirm the presence of free CHR molecules not involved on the holding together the CHR-OTX008-SBECD macrostructure (Figure 6.).

### 2.7. Thermal analysis

The thermal analysis of all components and their binary and ternary mixtures revealed that properties of all components were affected when mixed (Table S1. and Figures S13-S14.). In the ternary mixture, all components degrade together with a peak at 300 ^°^C indicating the formation of a unique macrostructure while those CHR [18] molecules accomodated in the free volume of the macrostructure kept their thermal identity with melting point at 290 ^°^C.

### 2.8. Computational studies on CHR-OTX008-SBECD interactions

Initial supramolecular structures for the interactions between CHR, OTX008 and SBECD were optimized with the GFN2-xtb quantum semi-empirical method. This method has been previously shown to produce good quality geometries for a range of species, from small organics to polymers.

To gain a better understanding of the relevant interactions, binding free energies have been calculated with DFT calculations (RI-PBE_D3BJ/def2-tvzp level of theory). All starting geometries were taken from the optimized GFN2-xtb structures.

The calculated geometries of optimized binding interactions between all molecular components (OTX008, CHR, SBE and CD) are shown in Figure 7. with calculated free energies in reported in Table 2.

**Figure 7.**
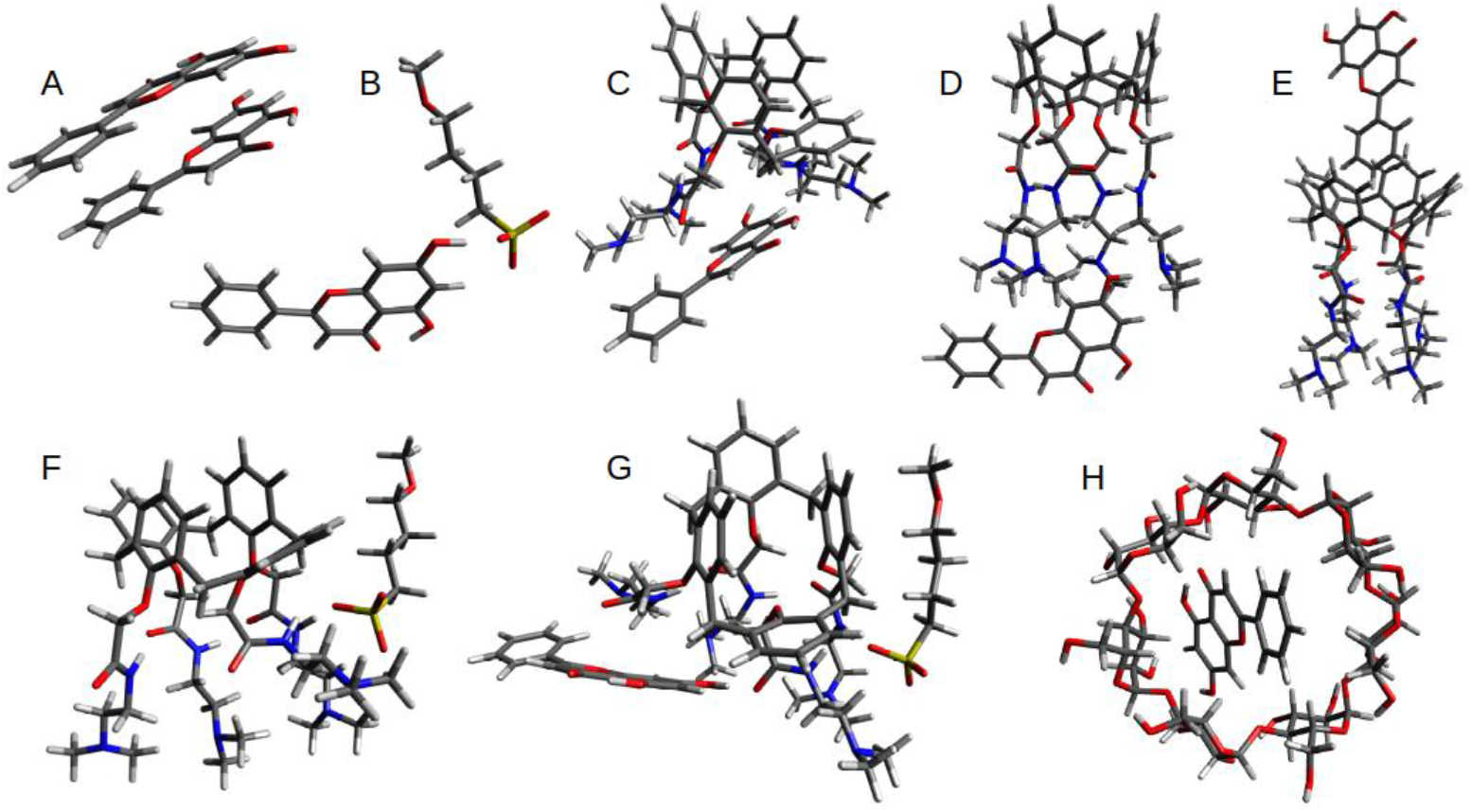
The figure shows the DFT optimized structures evaluated in this study, see Table 2 for calculated binding energies. Structure A shows the preferential CHR-CHR dimer alignment, with a binding energy of −14.0 kcal mol^-1^, alternative stacked orientations were within 2 kcal mol^-1^, suggesting multiple alignment possibilities. The hydrogen bonded dimer adducted was calculated to be 9 kcal mol^-1^ less favorable. Structure B highlights the CHR-SBE interaction, with a strong hydrogen bond between the -OH group of the CHR molecule and SO_3_^-^ group of the SBE arm. Structures C-E show the possible interactions between CHR and OTX008; with hydrogen bonding between the CHR hydroxyl group and the carbonyl on the OTX008 unit being preferential. Structure F shows the most favored interaction between the amine groups of the OTX008 unit and the SBE arm. Structure G highlights the tertiary adduct involving CHR-OTX008-SBE, the structure shows a cooperative effect where the OTX008-SBE interaction (R-NH---SO3-R) facilitates the OTX008-CHR interaction (R-CO---HO-R). This compensates for the increased entropic contribution of the tertiary complex. Structure H shows the host-guest binding of the CHR in the CD cavity.

**Table 2.** Binding free energies for each interaction. See Figure 7. for the structures.

| Structure | Interaction | G<sub>solv</sub> kcal/mol |
| --- | --- | --- |
| A | CHR dimer | -14.0 |
| B | SBECD-CHR (arm) | -6.2 |
| C | OTX008-CHR (carbonyl) | -12.6 |
| D | OTX008-CHR (amine) | -10.8 |
| E | OTX008-CHR (calixarene) | -2.7 |
| F | SBECD-OTX008 | +0.9 |
| G | SBECD-OTX008-CHR | -8.6 |
| H | SBECD-CHR (cavity) | -4.5 |

(CHR- Chrysin, SBECD- Sulfobutylated β-cyclodextrin sodium salt).

Given the size and conformation flexibility of the SBECD molecule, full DFT calculations would prove computational too expensive while also have a significant level of uncertainty with the results. A model system for the binding interaction between SBECD, CHR and OTX008 was chosen to focus on the SO_3_^-^ interactions as these showed to the preferential in the GFN2-xtb optimization. The GFN2-xtb optimization interactions between SBECD and CHR, and OTX008 and SBECD are also presented in Figure S15.

The relative binding energies (Table 2.) clearly highlight the preferential binding of the OTX008-CHR interaction via the carbonyl of the amide arms. There is also a reasonably strong interaction between the SBECD and CHR molecules both with the arms and the cyclodextrin cavity of SBECD. Interestingly the SBECD-OTX008 interaction is not favorable (+0.9 kcal mol^-1^). This agrees with the reduction in binding energy on the combination of SBECD-OTX008-CHR compared to the initial OTX008-CHR interaction. The formation of CHR dimers is also preferential.

### 2.9. Cell viability

Exposure to CHR, SBECD, their combinations with CHR, DMSO, or M did not impair H9c2 cell survival in either normal glucose (NG) or high glucose (HG) mediums. Furthermore, all tested concentrations of OTX008 (2.5, 1.25, 0.75 µM), whether alone or in combination with SBECD (OTX008-SBECD), as well as the mixture CHR-SBECD with OTX008 (CHR-OTX008-SBECD), did not diminish cell viability under NG conditions, as depicted in Figure 8A. These concentrations were also non-toxic in HG medium, as shown in Figure 8B. On the contrary, they lead to a significant improvement in cell viability on H9c2 cells exposed to HG. Notably, OTX008 alone and its combinations with SBECD markedly enhanced cell viability at all three tested doses (as seen in Figures 8C, D, E). The ternary mixtures of CHR-OTX008-SBECD began to show effectiveness at 1.25 µM OTX008 concentration, according to Figure 8D. It is particularly interesting that the combination of OTX008-SBECD, or CHR-SBECD-OTX008 resulted in the most pronounced increase in cell viability at the highest OTX008 dose of 2.5 µM, as illustrated in Figure 8C.

**Figure 8.**
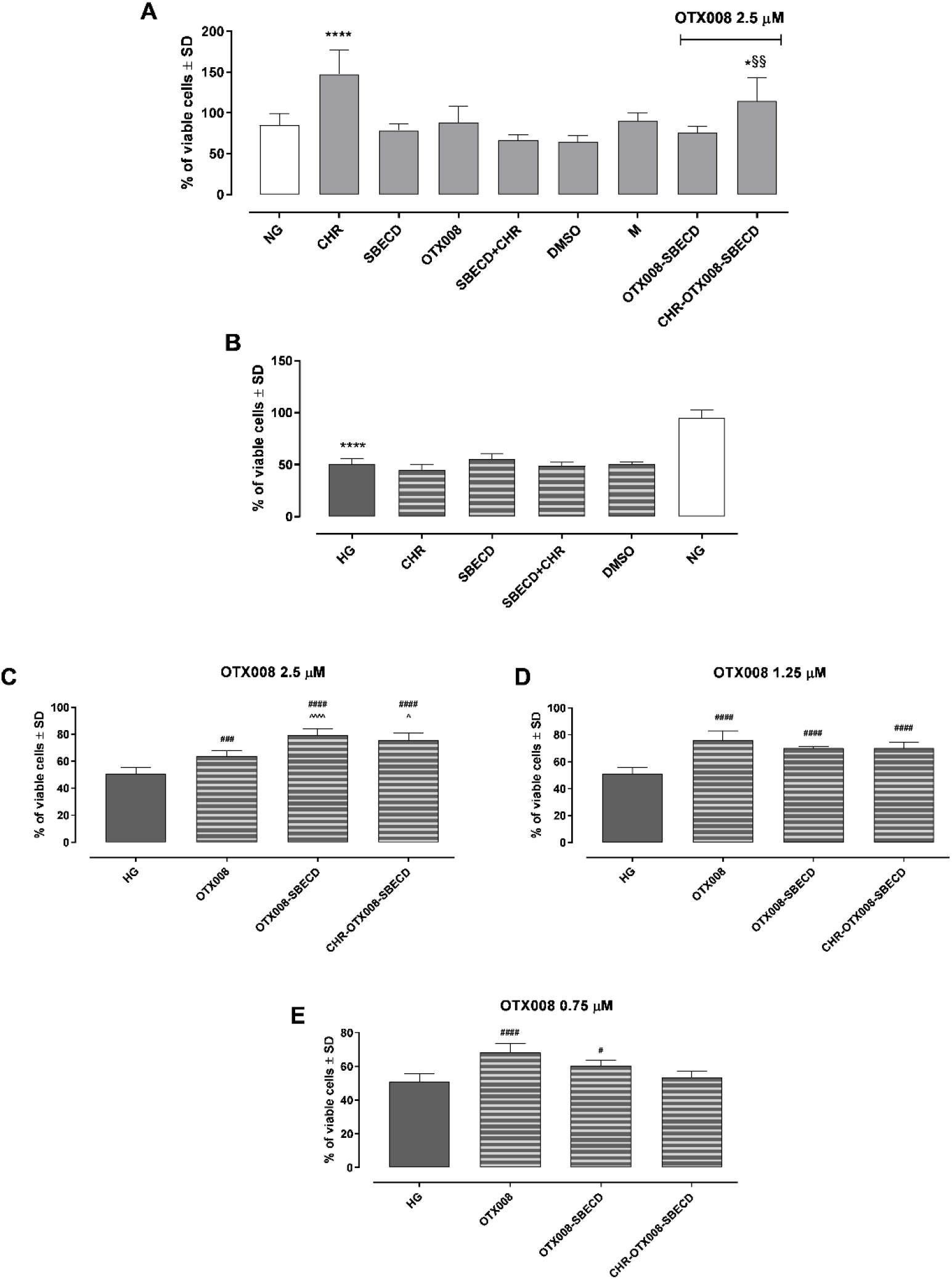
H9c2 cell viability after the treatment with the compounds in NG (A) and HG (B) conditions (where NG group is reported as control). In NG cells, OTX008 was tested at the maximum dose of 2.5 µM. In HG cells, OTX008 was tested at the doses of 2.5-1.25-0.75 µM (C, D, E respectively). Cell viability was determined by MTT assay and reported by % of cell viability ± SD. NG = 5.5 mM D-glucose; HG = 33 mM D-glucose; CHR = CHR 0.399 mg/ml; SBECD = Sulfobutylated-β-cyclodextrin 7.3 m/m%; SBECD+CHR = SBECD+0.095 mg/ml CHR; DMSO = dimethyl sulfoxide 2.5%; M = mannitol 27.5 mM; OTX008 = OTX008 (2.5-1.25-0.75 µM); OTX008-SBECD = OTX(2.5-1.25-0.75 µM)-SBECD; CHR-OTX008-SBECD = CHR (0.324 mg/ml)-OTX(2.5-1.25-0.75 µM)-SBECD. *P < 0.05, **** P < 0.0001 vs NG; # P < 0.05, ### P < 0.001, #### P < 0.0001 vs HG; ^ P < 0.05, ^^^^P < 0.0001 vs OTX008; §§ P < 0.01 vs OTX008-SBECD.

### 2.10. Reactive Oxygen Species (ROS)

Exposure to CHR, SBECD, their combinations with CHR, DMSO, or M did not affect ROS levels in either NG or HG conditions (Figure 9A). As expected, H9c2 cells grpwth in HG medium showed a marked increase in ROS intracellular levels compared to NG group (P < 0.001) (Figure 9B). Furthermore, all tested concentrations of OTX008 (2.5, 1.25, 0.75 µM), whether alone or in combination with SBECD (OTX008-SBECD), significantly reduced ROS levels in HG conditions (Figure 9C,D,E). Notably, the ternary mixtures of CHR-OTX008-SBECD began to show effectiveness at 1.25 µM OTX008 concentration, resulting in the most pronounced decrement of intracellular ROS levels at the highest OTX008 dose of 2.5 µM, as illustrated in Figure 9C.

**Figure 9.**
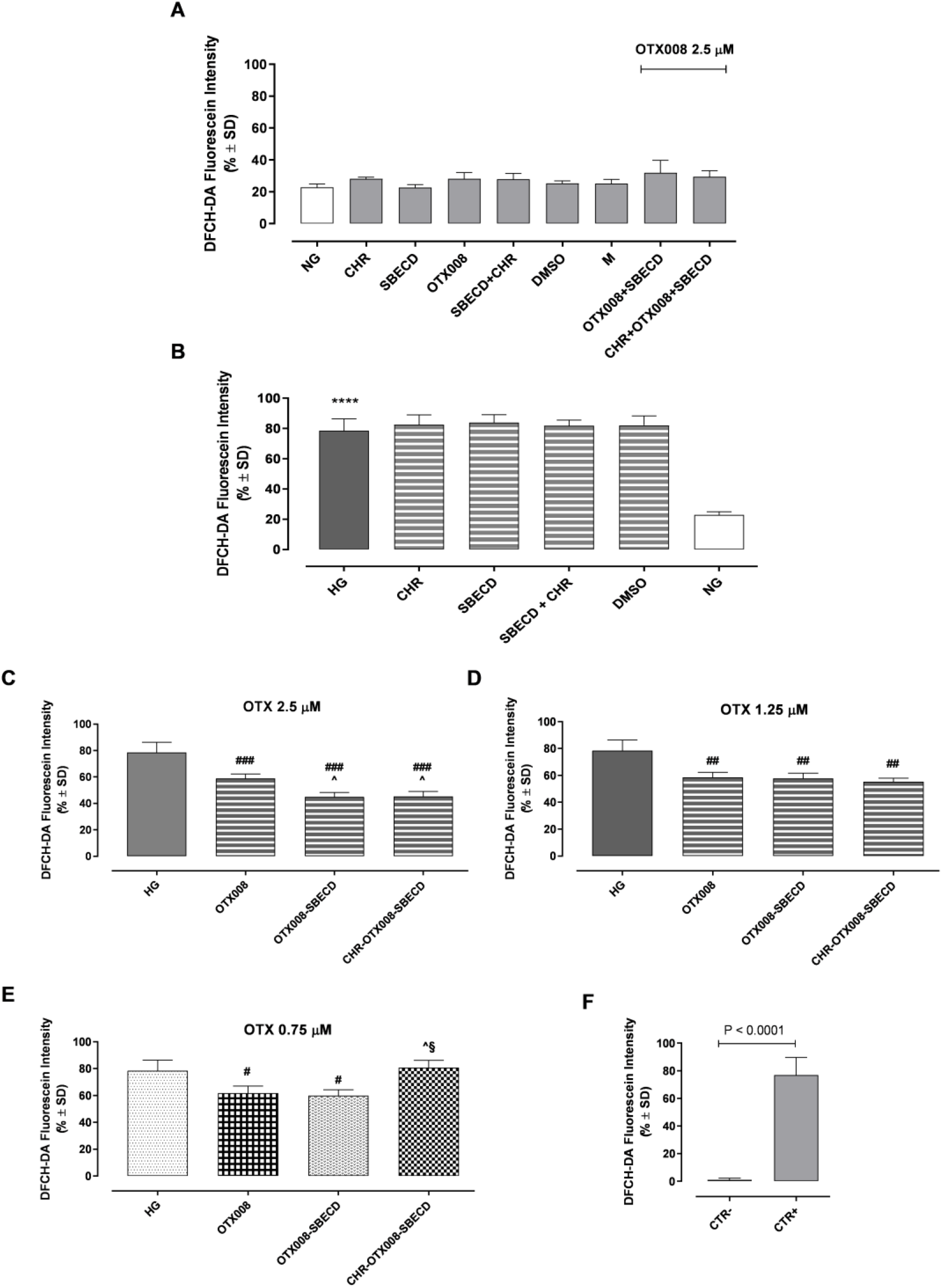
(A) 2′,7′-dichlorodihydrofluorescein diacetate (DCFH-DA) levels in NG or (B) HG medium (where NG is reported as control). In NG cells, OTX008 was tested at the maximum dose of 2.5 µM. In HG cells, OTX008 was tested at the doses of 2.5-1.25-0.75 µM (C, D, E respectively); (F) CTR– = negative control (5% FBS without DCFH-DA); CTR+= positive control (H_2_O_2_ 100 µM). DCFH-DA probe was used to assessed ROS levels, quantified on a flow cytometer (Figure S15), and reported as DCFH-DA fluorescein intensity ± SD. NG = 5.5 mM D-glucose; HG = 33 mM D-glucose; CHR = CHR 0.399 mg/ml; SBECD = Sulfobutylated-β-cyclodextrin 7.3 m/m%; SBECD+CHR = SBECD+0.095 mg/ml CHR; DMSO = dimethyl sulfoxide 2.5%; M = mannitol 27.5 mM; OTX008 = OTX008 (2.5-1.25-0.75 µM); OTX008-SBECD = OTX(2.5-1.25-0.75 µM)-SBECD; CHR-OTX008-SBECD = CHR (0.324 mg/ml)-OTX(2.5-1.25-0.75 µM)-SBECD. **** P < 0.0001 vs NG; # P < 0.05, ## P < 0.01, ### P < 0.001 vs HG; ^ P < 0.05 vs OTX008; § P < 0.01 vs OTX008-SBECD.

## 3. Discussion

Chronic hyperglycemia-induced fibrosis affects vascular structures, leading to diabetic complications like retinopathy and liver fibrosis [19]. This fibrogenic process involves inflammatory mediators, cytokines, and growth factors, especially TGF-β. This leads to increased extracellular matrix deposition. Past research on the flavonoid CHR in rodents showed its anti-fibrotic effects. CHR counteracts fibrosis by inhibiting hepatic cell activation via the TGF-β1/Smad pathway [20]. Additionally, CHR adjusts extracellular matrix dynamics and reduces collagen synthesis [11].

Recent studies highlight Gal-1 protein as a therapeutic target for diabetic fibrosis. Elevated Gal-1 levels were found in diabetic mice kidneys, contributing to kidney fibrogenesis [21,22]. Additionally, OTX008 has proven effective in preventing Gal-1 accumulation and countered the effects of TGF-β on ARPE-19 cells in high glucose situations [23].

In this study, we suggest the concurrent administration of CHR and OTX008 as a strategy to combat fibrosis. However, this approach requires an appropriate delivery system to facilitate their combined solubilization and delivery. SBECD was chosen as the preferred carrier due to its established safety for parenteral use, superior solubilizing capabilities, and polyanionic character [24–26]. Previous research has demonstrated the successful solubilization of CHR using SBECD [5]. The potential interaction between SBECD and OTX008 was inferred from their chemical structures. Subsequent solubilization tests and chemico-physical examinations confirmed that SBECD, OTX008, and CHR interact to create a ternary complex, with initial findings indicating that SBECD is capable of dissolving OTX008. Phase solubility tests of CHR clearly showed the difference between the solubilization mechanisms and the structure of the formed complexes in the binary CHR-SBECD and ternary CHR-OTX008-SBECD complexes. The positively skewed curve of the ternary complexes points to the fact, that bigger molecular associates were formed and increased the solubility of CHR compared to SBECD. This phenomenon is well-known in the case of cyclodextrins [27], however in calixarene-cyclodextrin macrostructures it is less studied. The solubility of CHR is pH-dependent, in the range of pH 6.20-9.40 the two OH groups of the molecule are ionizable and the two ionizations take place forming a dianion [28]. It is in accordance with our results, in the presence of SBECD increasing the pH from 3.65 to 9.95 resulted approximately a twofold increase in CHR solubility. SBECD-CHR host-guest interaction is favorable from computational calculations (−4.5 kcal mol^-1^), supported also by the NMR data, with CHR accommodated inside the toroid cavity of the cyclodextrin ring. Molecular simulation revealed another interesting mechanism in the interaction of CHR and SBECD. The OH groups of CHR chemically interacts with the sulfobutyl ether (SBE) pendant groups, which is not the conventional host-guest mechanism of cyclodextrins but can contribute to the solubilization of CHR. SBECD has an average of six SBE arms, thus it can amplify the number of interactions with CHR molecules, beside the host-guest interaction. The solubilization of OTX008 with SBECD resulted also an alkaline solution with a pH of 9.36, however increased the CHR solubility fourfold, showing, that the interactions with OTX008 was required for the significant CHR solubilization. In the ternary complex of CHR-OTX008-SBECD the NMR signals of CHR’s OHs are significantly changed, confirming the involvement of OH groups in the formation of ternary complexes. The signals of the amide group and aromatic rings of OTX008 are also modified showing involvement on the interaction in the ternary complex. Indeed, the most preferential interactions are on the carbonyl and amine groups (−12.6 kcal mol^-1^ and −10.8 kcal mol^-1^ respectively) of OTX008 with CHR.

The presented geometries and binding energies suggest that the experimentally enhanced entrapment of CHR with an OTX008-SBECD mixture is additive in nature and the higher solubility observed can be justified by a combination of these mechanisms, leading to a supramolecular structure. The SBECD is able to bind at least 6 CHR molecules (one on each arm) with the potential for multiple CHR per SO_3_^-^ moiety. Subsequently the addition of OTX008 allows for 1:1 CHR-OTX008 adducts to be formed. The CHR is preferentially located within the amide arms (allowing for additional secondary interactions) and optimization an external amide interaction proves less stable, suggesting the 1:1 complex most favored.

The hypothesis here is that the OH groups of CHR molecules located in position 7 (Figure 6A.) are involved in hydrogen bonding with OTX008 molecules (CONH) and/or in interactions with the tails of SEBCD to produce a supramolecular structure; the higher solubilization of the CHR in the ternary structure is due to the cooperation of three effects: i) electrostatic bonding of CHR with OTX008 and SBECD molecules acting as bridge, ii) inclusion of CHR molecules in the SBECD toroid and calixarene structures and iii) entrapment of CHR molecules in the free volume of the new supramolecular structure, facilitated by the strong CHR-CHR binding energy (Figure 10.). This high solubilization of CHR is strictly associated to this specific ternary system.

**Figure 10.**
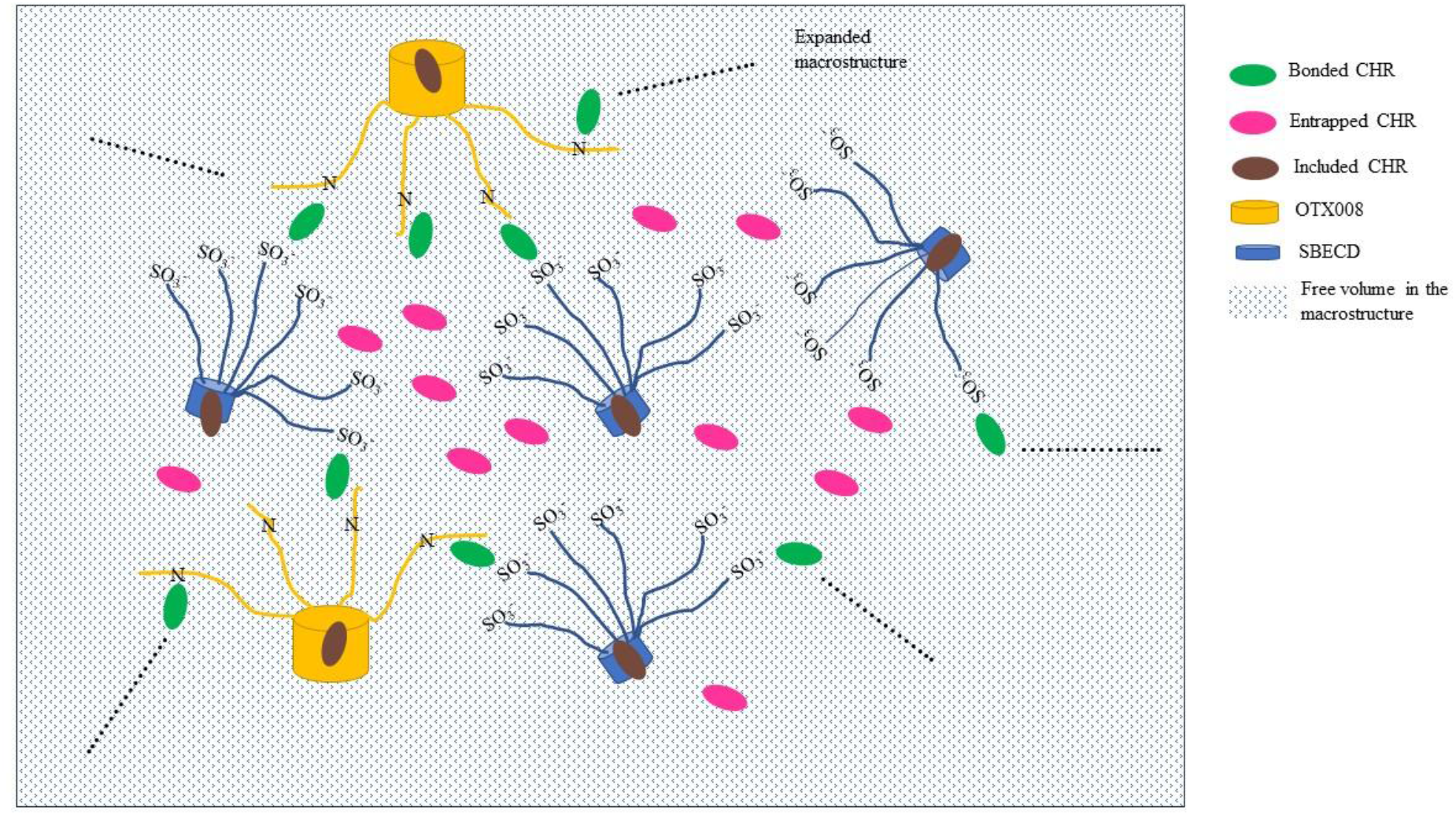
Schematic representation of CHR-OTX008-SBECD macrostructure with CHR molecules making the link between SBECD and OTX008 molecules (Green discs), dispersed in the free volume created in the macrostructure (Purple discs) and trapped in the CD toroid or calixarene structure (Brown discs). The synergy of the three situations gives the enhancement of the CHR solubility in the ternary mixture.

Based on the phase solubility data and DLS results large molecular associations are formed in the solution of OTX008 and SBECD (1002 nm, 86.8% intensity), which further increased after the solubilization of CHR (2698 nm, 75.2% intensity). Between the molecules, which forms the supramolecular associations free volume takes place [29,30] where CHR can be accumulated by the supramolecular structure. The special structure of the two association builder macrocycles OTX008 and SBECD helps the accumulation of CHR between macro chains or side groups. The structure supports the formation of the favored supramolecular carrier by the binding free energies, complexation, and free volumes.

An attempted scheme with the proposed supramolecular structure is reported in Figure 10.

Finally, the biocompatibility of the complexes was tested on H9c2 (2-1) cells under normal and hyperglycemic conditions. The tested compounds alone, binary, and ternary complexes were not toxic on cells in NG or HG medium. Moreover, the binary and ternary complexes of OTX008 prevented the toxic effects of HG medium on H9c2 cells, especially at the maximum dose of OTX008 (2.5 µM) was the most effective. The same trend was shown by the binary and ternary complexes of OTX008 in reducing intracellular ROS levels observed in H9c2 cells exposed to HG condition. Building on our prior research that highlighted CHR’s antifibrotic capabilities [11,20] and OTX008’s intervention in the profibrotic Gal-1/TGF-β pathway in a hyperglycemic environment [21], this newly devised drug delivery system CHR-OTX008-SBECD emerges as a promising candidate for modulating fibrotic progression associated with chronic diabetes. To this regard, its anti-fibrotic properties have been verified in H9c2 exposed to hyperglycemic conditions, while its safety and efficacy have been tested intraperitoneally in adult CD1 male mice with chronic diabetes. In this in vivo experimental setting, CHR-OTX008-SBECD lead to a significant reduction of cardiac fibrosis markers and cardiac tissue remodeling, without any toxicity, by improving both CHR and OTX008 solubility in water and bioavailability [31]. Further detailed in vitro and in vivo studies are essential to discern the specific cellular and molecular mechanisms driving its therapeutic potential.

## 4. Materials and Methods

### 4.1. Materials

OTX008, known as Calixarene 0118, was obtained from Selleck Chemicals GmbH. The sulfobutylated β-cyclodextrin sodium salt (SBECD) with a degree of substitution (DS) around 6, was procured from Cyclolab Ltd. in Budapest, Hungary. CHR, chemically identified as 5,7-Dihydroxyflavone, was purchased from Alfa Aesar (by ThermoFisher Scientific in Kandel, Germany), while all other chemical reagents were supplied by Sigma.

### 4.2. Methods

#### 4.2.1. Solubilization studies

##### 4.2.1.1. OTX008 solubilization with SBECD (Binary systems)

SBECD solutions with increasing concentrations were prepared using ultrapure water (Millipore Direct-Q 5UV system, Merck Millipore, Burlington, MA, USA) and the same amounts of OTX008 were tested to dissolve in them to reveal the solubilization ability of SBECD on OTX008. OTX008 was clearly dissolved in the 7.3 m/m% SBECD solution at a 0.75 mg/ml final concentration to get OTX008-SBECD solution.

##### 4.2.1.2 CHR solubilization in OTX008-SBECD solution (Ternary systems)

CHR was added in excess to solutions of OTX008-SBECD, prepared as per method 4.3.1, and agitated for 72 hours at room temperature in closed vials (n=3). Post-incubation, the mixtures were centrifuged at 11,000 rpm for 10 minutes. The resulting clear supernatants were then separated, and the solubilized CHR concentration was determined using a UV spectrophotometer (Shimadzu UV-1900) at a wavelength of 270 nm. CHR solubilization was similarly conducted in the solution of 7.3 m/m% SBECD in the absence of OTX008, following the same incubation and preparation process.

The clarified CHR solutions were subsequently frozen at −110 °C and lyophilized using a ScanVac CoolSafe freeze dryer (Labogene, Allerød, Denmark). The resulting complexes were preserved at −20 °C for further experiments.

##### 4.2.1.3 Phase-solubility study

Solutions of 7.3 m/m% SBECD, and OTX008-SBECD, were made and then diluted in a 96-well plate with ultrapure water. A surplus of CHR was added to each well. The plates were then sealed and agitated for 72 hours at room temperature. After shaking, the samples were passed through a MultiScreen Solvinert 96 Well Filter Plate with a 0.45 µm pore size, Poly(tetrafluoroethene) membrane (PTFE) (Merck Millipore Ltd., Tullagreen, Ireland), using a MultiScreen Resist vacuum manifold (EMD Millipore corporation, Burlington, MA, USA). The filtered, clear supernatants were transferred to a Greiner UV-Star® 96 well plate and the absorbance was measured at 270 nm using a Thermo Fisher Multiskan Go microplate reader (Thermo Fisher, Waltham, MA, USA).

##### 4.2.1.4 pH-dependent CHR solubility determination

SBECD solutions at a concentration of 7.3 m/m% were prepared using a sodium acetate-acetic acid buffer at various pH levels (pH= 3.65; 5.75; 7.45; 9.95) and dispensed into a 96-well plate. An excess of CHR was then added to each well, the plate was sealed, and the solubility of CHR was assessed as described in section 4.3.3.

In a different experimental setup, the pH levels of the 7.3 m/m% SBECD and OTX008-SBECD solutions were adjusted to pH 8.64 with a 0.2 M sodium acetate solution and to pH 3.65 with 0.1 M HCl, respectively. Following these pH adjustments, the solubilization test for CHR was carried out as previously described.

### 4.3. Scanning Electron Microscopy (SEM) Analysis

The lyophilized samples of complexes were crushed and mounted on a fixture with graphite-containing, double-sided adhesive tape. The pure raw materials were used without crushing. The surface of the samples was not coated with gold before the SEM examination. A vacuum and low accelerating voltage of 5-15 kV were used during the investigation by a Hitachi Tabletop microscope (TM3030 Plus, Hitachi High-Technologies Corporation, Tokyo, Japan).

### 4.4. Size distribution measurement of the cyclodextrin complexes with dynamic light scattering (DLS)

The particle size distribution in solutions of 7.3 m/m% SBECD, and OTX008-SBECD, both with CHR complexed after the phase solubility tests and without CHR, was measured using a Malvern Nano-ZS Zetasizer (Malvern Instruments, Malvern, UK).

### 4.5. Nuclear Magnetic Resonance (NMR) studies

^1^H and NOESY NMR characterisation of CHR, SBECD, OTX008, binary systems CHR-SBECD, OTX008-SBECD, and ternary system CHR-OTX008-SBECD were performed using a Bruker Ascend 400 MHz spectromenter with a double resonance broad band probe, BBFO model, and the spectra recorded at ambient temperature in dimethylsulphoxide (DMSO-d_6_) and/or deuterated water (D_2_O) depending on the solubility of the materials (Typical solutions of 10 mg/mL). Lyofilised mixtures CHR-SBECD, OTX008-SBECD, and CHR-OTX008-SBECD were used as described 4.3.1 and 4.3.2. The solvent peaks were referenced to 2.5 ppm (DMSO-d_6_) and 4.7 ppm (HDO, H_2_O). Peak multiplicities were described as follows: singlet (s), multiplet (m), and broad (br).

### 4.6. Differential Scanning Calorimetry (DSC) studies

The melting temperature (T_m_) and degradation temperature (T_d_) of all components and their binary and ternary mixtures were determined by Differential Scanning Calorimetry (DSC; Mettler Toledo, DSC 30 STAR System) at the heating rate of 10 ^°^C min^-1^ and under an inert N_2_ atmosphere.

### 4.7. Computational studies

All calculations were undertaken using the Orca 4.2.1 software package [32]. Initial geometry optimization for CHR, OTX008 and SBECD and all interaction variants were conducted with GFN2-xtb semi-empirical quantum mechanical method [33]. The method has been shown to be robust for large scale calculations (up to a few thousand atoms) of organic, organometallic and biochemical systems. The inclusion of the D4 dispersion term makes GFN2-xtb ideal for studying structures involving non-covalent interactions [34].

To gain a more accurate energetic understanding of the molecular interactions the structures were re-optimized with unconstrained Density Functional Theory (DFT) calculations, using the RI-PBE(D3)/def2-svp level of theory [35–38], with subsequent solvated single point energy calculations at the RI-PBE(D3)/def2-tzvp level of theory [35– 38] with the continuum solvent polarized method (cpcm), [39] simulating a water environment (epsilon = 80.3). Analytical frequency calculations on the gas phase geometries were performed to account for enthalpic and entropic contributions to the free energy term, as well as to confirm all intermediates are true minima on the potential energy surface. Due to the scaling limitations of DFT, and focus on the molecular interactions, the SBECD molecule was approximated by a single SBE arm when calculating the binding free energy for these interactions. A concentration-induced free-energy shift of *R T* ln*V*_M_=1.89 kcal mol^−1^ (V_M_: molar volume of an ideal gas, *T*=298 K) has been included to account from the shift from gas (1 atm) to solution (1 mol dm^-3^) phase.

### 4.8. H9c2 cell culture

Embryonic rat cardiac H9c2 (2-1) cells (ECACC, United Kingdom) were cultured in Dulbecco’s modified Eagle’s medium (DMEM; Aurogene, Italy), containing 5.5 mM d-glucose and supplemented with 10% heat inactivated fetal bovine serum (FBS; AU-S181H Aurogene, Italy), 1% L-Glutamine (L-Glu; AU-X0550 Aurogene, Italy) and 1% penicillin/streptomycin solution (P/S; AU-L0022 Aurogene, Italy), at 37°C under an atmosphere of 5% CO2.

Reached an 80% confluence, H9c2 cells were trypsinized, seeded at a specific cell density for each assay and then exposed to NG, high glucose (HG; 33 mM d-glucose) or NG + 27.5 mM mannitol (M; as osmotic control) for 48 hours [40]. Cells were then treated for 6 days [41] in NG or HG medium with the following substances:

- CHR 0.399 mg/ml (CHR), dissolved in NaCl;
- SBECD 7.3 m/m%, dissolved in NaCl;
- Binary system SBECD+0.095 mg/ml CHR (SBECD+CHR), dissolved in NaCl;
- DMSO 2.5% as vehicle of OTX008;
- OTX008 (0.75-1.25-2.50 µM);
- Binary system OTX008 (2.5-1.25-0.75 µM)-SBECD (OTX008-SBECD), dissolved in NaCl;
- Ternary system CHR (0.324 mg/ml)-OTX008 (2.5-1.25-0.75 µM)-SBECD (CHR-OTX008-SBECD), dissolved in NaCl.

Three independent experiments were done, each performed in triplicates (N = 9).

### 4.9. Cell viability assay

H9c2 cells were plated at a density of 1 x 10^4^ cells per well in 96-well plates [42], exposed to NG or HG medium for 48 hours and then treated as above described.

At the end of the stimulation period, 3-(4,5-Dimethylthiazol-2-yl)-2,5-Diphenyltetrazolium Bromide (MTT) solution (1:10 in culture medium, 300 µl/well) was added to each well, incubated for 4 h at 37°C and then removed. Each well was then washed for 20 min with isopropanol-HCl 0.2 N. Optical density (OD) values were measured at 570 nm using a 96-well plate reader (iMark, Bio-Rad Laboratories, Italy) [40].

### 4.10. ROS assessment

ROS levels were detected by the conversion of the fluorescent probe 2′,7′-dichlorodihydrofluorescein diacetate (DCFH-DA) to highly fluorescent dichlorofluorescein (DFC) diacetate within cells by ROS. H9c2 cells were seeded in 6-well plates (5 x 10^4^ cells/well) to NG or HG medium for 48 hours and then treated as above described. At the end of the stimulation period, cells were loaded with 20 µM DCFH-DA in medium with 5% FBS at 37°C for 30 min, then were trypsinized. Total intracellular ROS production was quantified on a Guava easyCyte flow cytometer (Merck Millipore, Burlington, MA, USA), following the manufacturer’s instructions. Both cell types were exposed to medium 5% FBS without DCFH-DA as negative control (CTR-) or incubated with H_2_O_2_ (100 µM) 30 min before trypsinization as a positive control (CTR+).

### 4.11. Statistical analysis

The results are reported as mean ± standard deviation (SD). Statistical significance was determined using one-way Analysis of Variance (ANOVA) followed by Tukey’s comparison test. A P-value less than 0.05 was considered significant to reject the null hypothesis.

## 5. Conclusions

Addressing fibrosis under hyperglycemic conditions remains a significant challenge, necessitating innovative approaches and therapeutic agents. In this study, we introduce a new ternary formulation and application of two potential drugs namely CHR and OTX008 with SBECD, each targeting distinct pharmacological pathways, within a Drug Delivery System (DDS) optimized for parenteral use. This marks the novel dual formulation of the calixarene derivative OTX008 and flavonoid CHR within a single DDS, where they uniquely and actively influence the structural formation of the resultant supramolecular ternary complex. Notably, OTX008 serves as a pivotal element of the ternary complex, both structurally and pharmacologically. Our findings suggest that the tailored ternary complex presents a novel promising solution for the integrated treatment of fibrosis in hyperglycemic scenarios and can open a new chapter in drug delivery of poorly water-soluble drugs.

## Supplementary Materials

The following supporting information can be downloaded at: www.mdpi.com/xxx/s1, Figure S1: Size distribution of the molecular associates of SBECD, OTX008-SBECD and CHR-OTX008-SBECD in water.; Figure S2: ^1^H NMR spectrum of OTX008 in DMSO-d_6_.; Figure S3: ^1^H NMR spectrum of SBECD in DMSO-d_6_.; Figure S4: ^1^H NMR spectrum of SBECD in D_2_O.; Figure S5: ^1^H NMR spectra of CHR-SBECD mixture, CHR and SBECD in DMSO-d_6._; Figure S6: ^1^H NMR spectra of CHR-SBECD mixture and SBECD in D_2_O.; Figures S7: NOESY NMR spectra of CHR, CHR-SBECD mixture and SBECD in DMSO-d_6_.; Figure S7a: Expanded area of the NOESY spectra Figure S7.; Figures S8: ^1^H NMR spectra of OTX008-SBECD mixture, OTX008 and SBECD in DMSO-d_6_.; Figures S9: NOESY spectra of OTX008-SBECD mixture, OTX008 and SBECD in DMSO-d_6_.; Figure S10: ^1^H NMR spectra of CHR-OTX008-SBECD mixture, OTX008, SBECD and CHR in DMSO-d_6_.; Figure S10a: Expanded area of ^1^H NMR spectra of CHR-OTX008-SBECD mixture, OTX008, SBECD and CHR in DMSO-d_6_.; Figure S10b: Expanded area of ^1^H NMR spectra of CHR-OTX008-SBECD mixture, OTX008, SBECD and CHR in DMSO-d_6_.; Figure S10c: Expanded area of ^1^H NMR spectra of CHR-OTX008-SBECD mixture, OTX008, SBECD and CHR in DMSO-d_6_.; Figure S11: Comparison of NOESY spectra of CHR, CHR-OTX008-SBECD mixture and binary CHR-SBECD mixture in DMSO-d_6_.; Figure S12: NOESY spectra CHR-OTX008-SBECD mixture and SBECD in D_2_O.; Figure S13: Zoomed view of DSC thermograms of CHR, OTX008, SBECD and CHR-OTX008-SBECD.; Figure S14: Comparison of zoomed areas of DSC thermograms of SBECD-OTX008 mixture, SBECD-CHR mixture and CHR-SBECD-OTX008 mixture.; Figure S15: Representative flow cytometer measures of total intracellular ROS levels assayed with DCFH-DA probe.; Figure S16: Interactions between SBECD and CHR, OTX008 and SBECD.; Table S1: Results of thermal analysis.

## Author Contributions

Conceptualization, A.He., E.D. A. Ha., M.D’A. and F.F; methodology, A.He., E. D., A. He., M. C. T., M.R., Á. R., J. V., F. F., CS. S., C. C. L., I. B., and I. Bu.; investigation, E.D., A.Ha., M. C. T., M.R., Á. R., J. V., F. F., CS. S., C. C. L., and I.Bu.; resources, A.He.; data curation, F.F.; writing—original draft preparation, A.He., E.D., A.Ha., F.F., Á.R., M.C.T., I.B., M.D’A and J.V.; writing—review and editing, A.He., E.D., A.Ha., and F.F.; All authors have read and agreed to the published version of the manuscript.

## Institutional Review Board Statement

Not applicable.

## Informed Consent Statement

Not applicable.

## Disclaimer/Publisher’s Note

The statements, opinions and data contained in all publications are solely those of the individual author(s) and contributor(s) and not of MDPI and/or the editor(s). MDPI and/or the editor(s) disclaim responsibility for any injury to people or property resulting from any ideas, methods, instructions or products referred to in the content.

