## Supplemental material for "Chrysin directing an enhanced solubility through the formation of a supramolecular cyclodextrin-calixarene drug delivery system: a potential strategy in antifibrotic diabetes therapeutics"

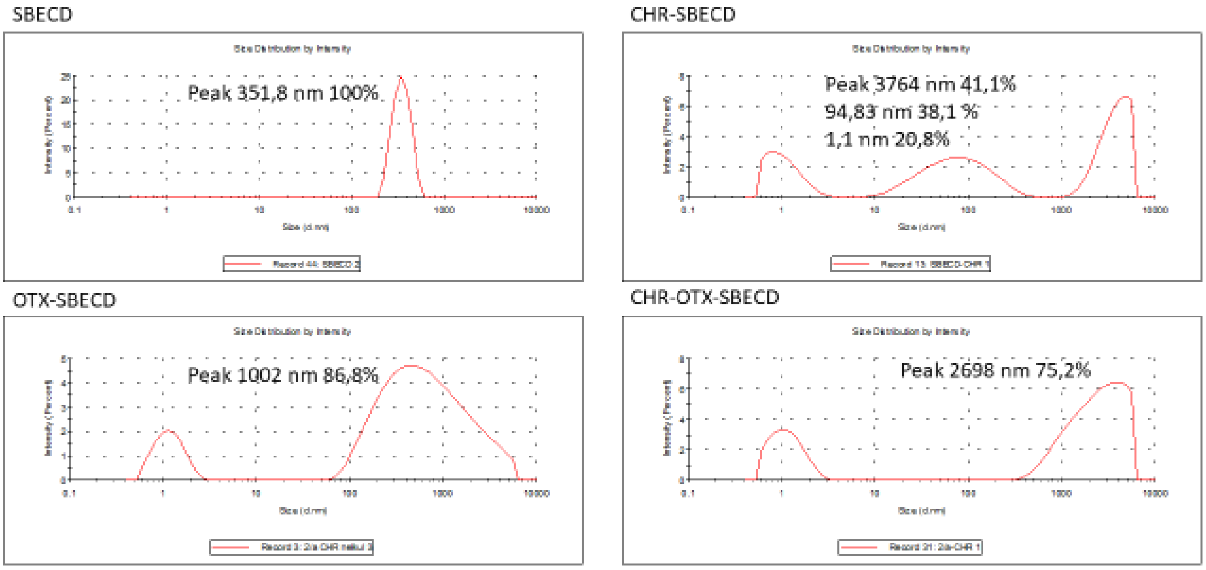


Figure S1. Size distribution of the molecular associates of SBECD, OTX008-SBECD and CHR-OTX008-SBECD in water. The associates were characterized by dinamyc light scattering. X-axis: size (nm); Y-axis: intensity distribution (%).

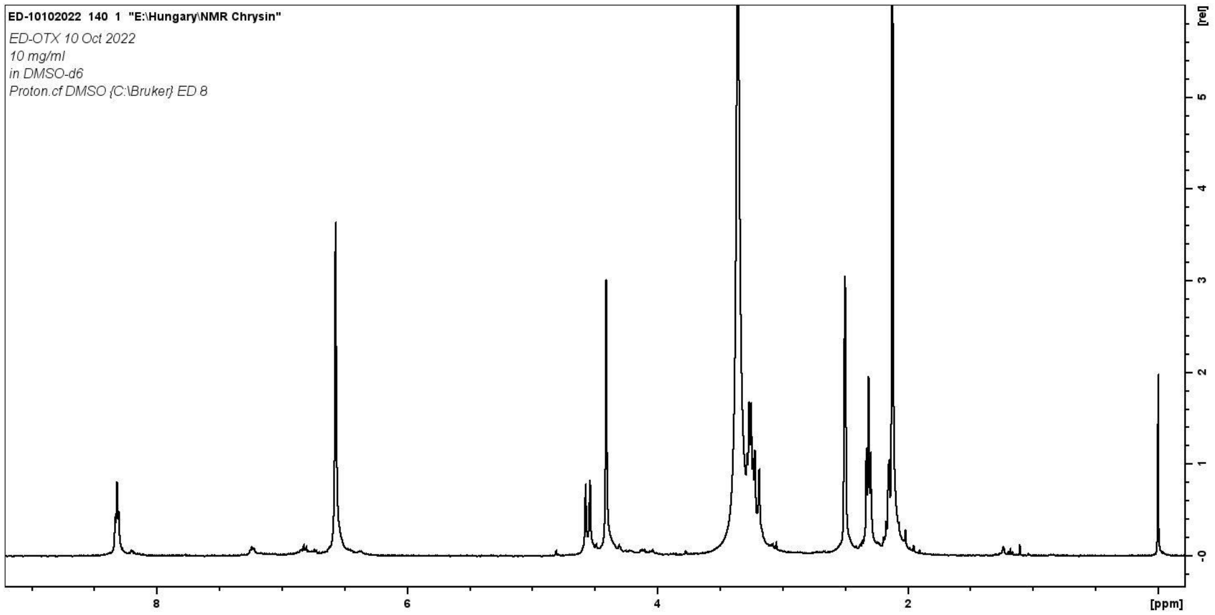


Figure S2. [1]H NMR spectrum of OTX008 in DMSO-$d_6$.

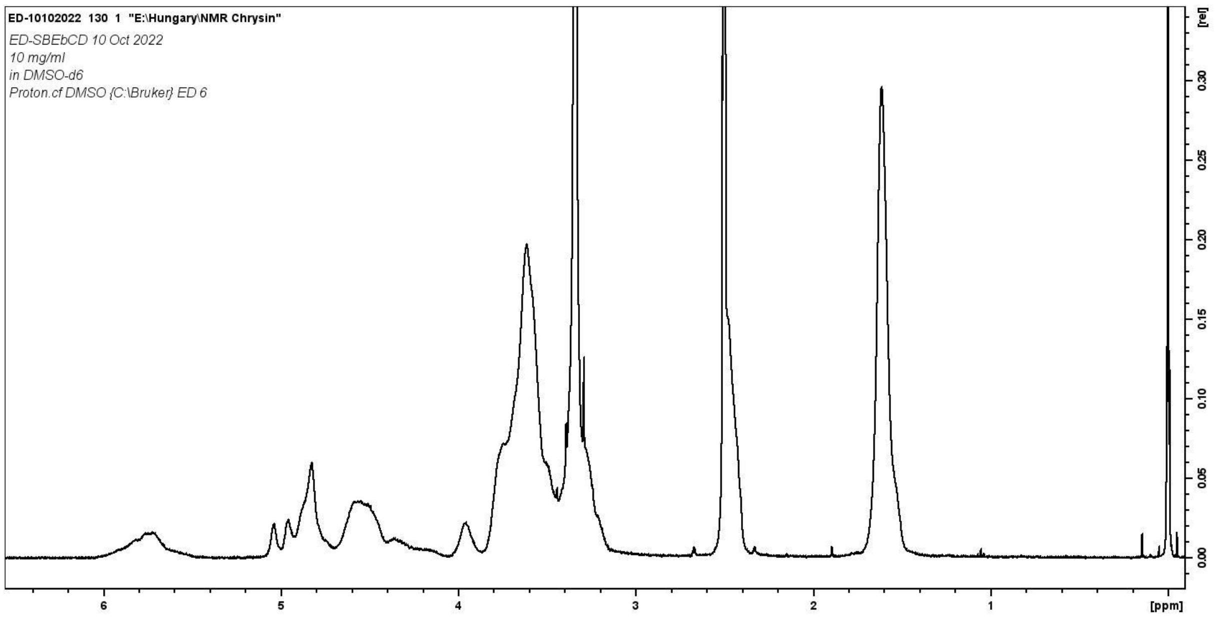


Figure S3. [1]H NMR spectrum of SBECD in DMSO-$d_6$.

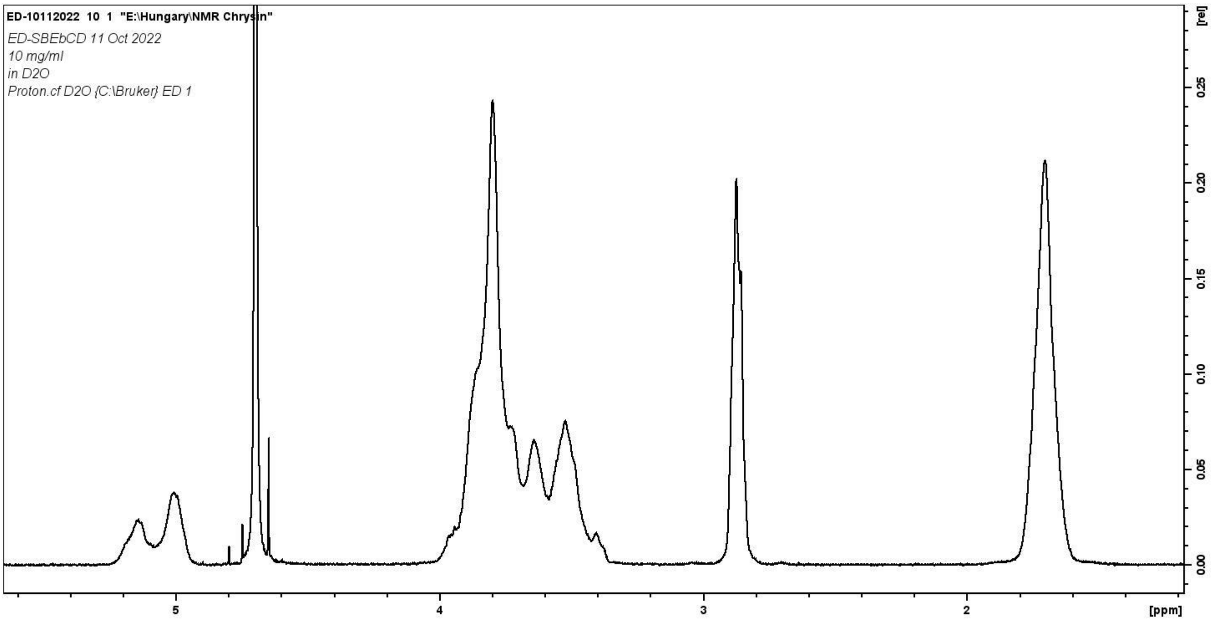


Figure S4. [1]H NMR spectrum of SBECD in $D_2O$.

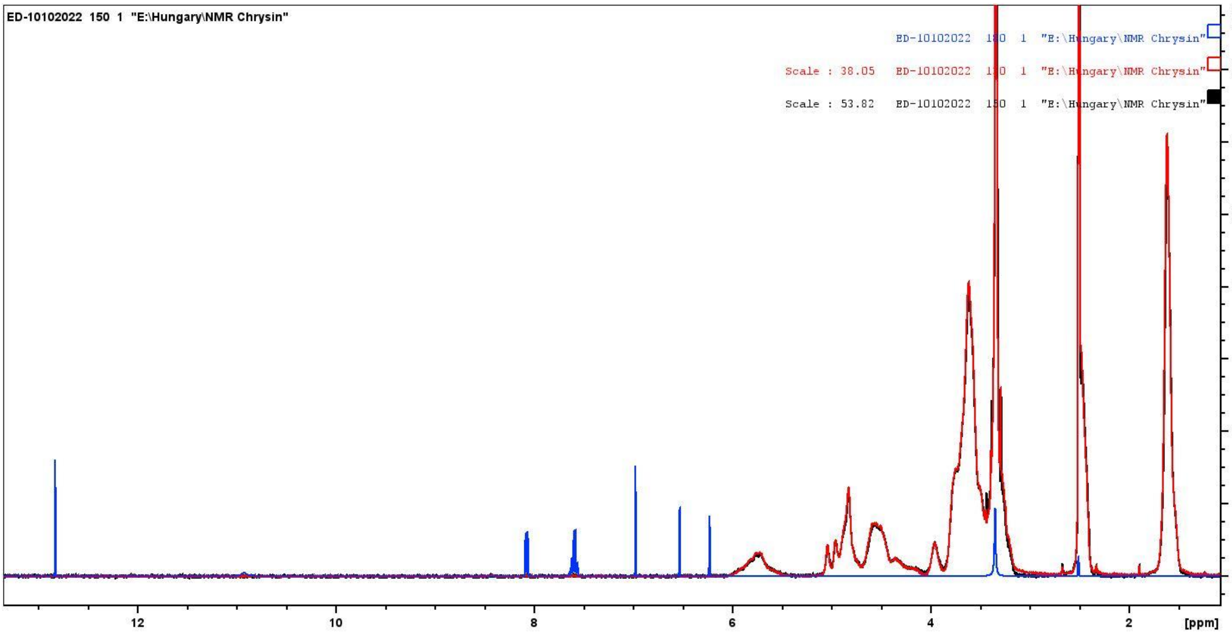


Figure S5. [1]H NMR spectra of CHR-SBECD mixture (black line), CHR (Blue line) and SBECD (Red line) in DMSO-$d_6$.

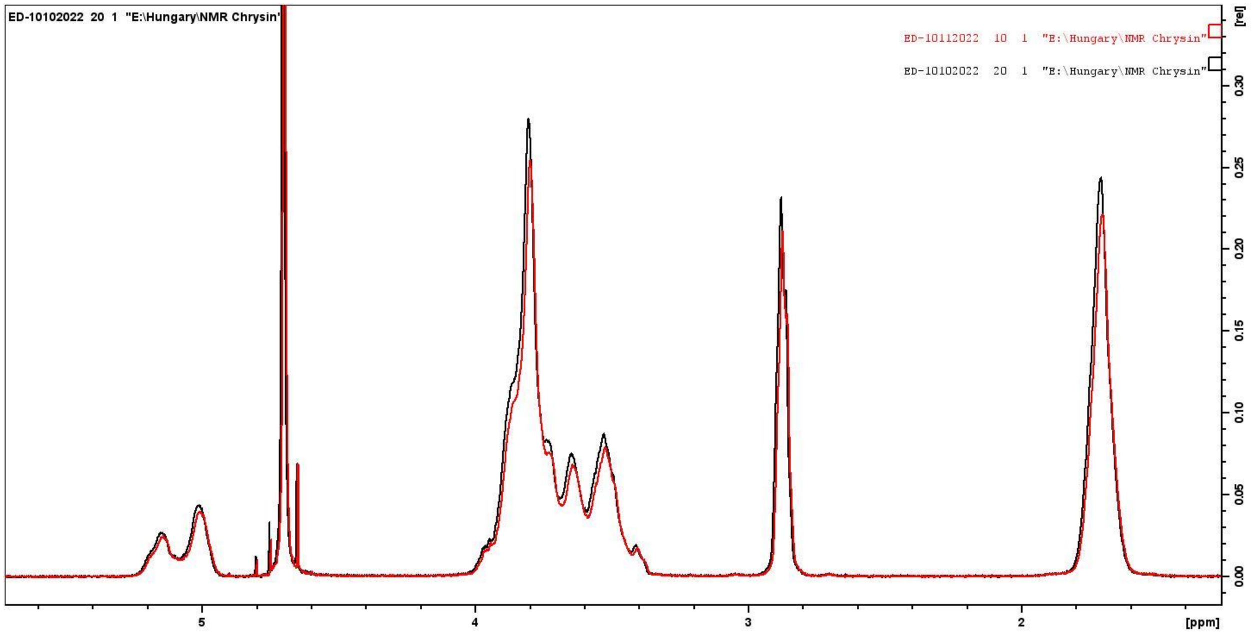


Figure S6. [1]H NMR spectra of CHR-SBECD mixture (black line) and SBECD (Red line) in $D_2O$.

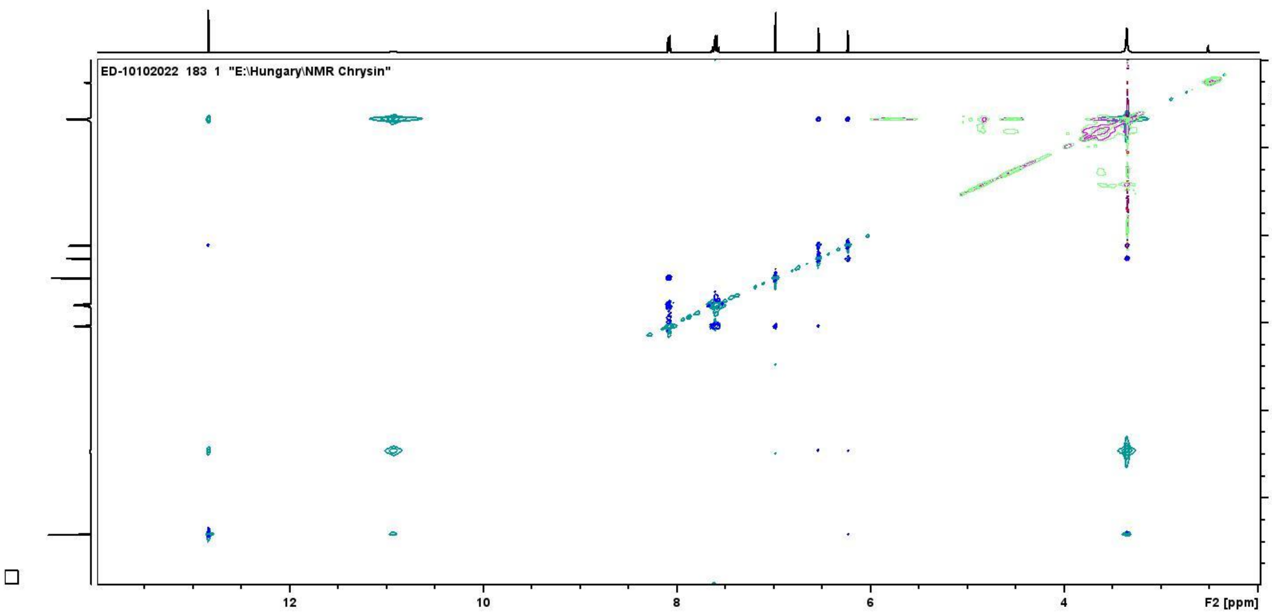


Figure S7. NOESY NMR spectra of CHR (Blue/Dark green line), CHR-SBECD mixture (Purple line) and SBECD (Light green line) in DMSO-$d_6$.

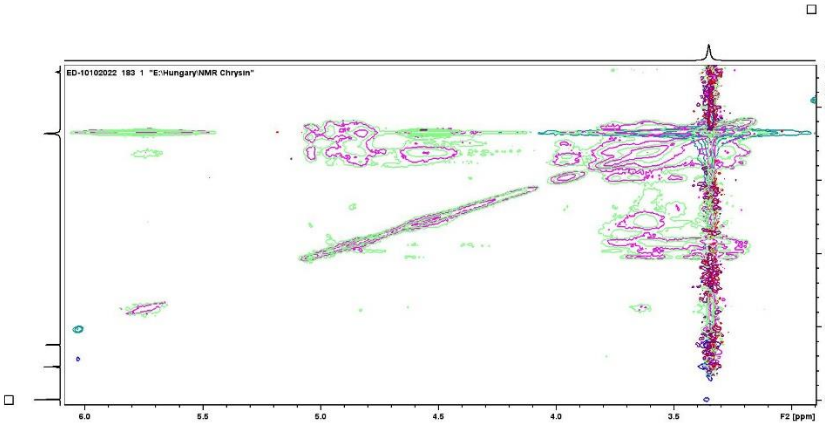


Figure S7a. Expanded area of the NOESY spectra Figure S7.

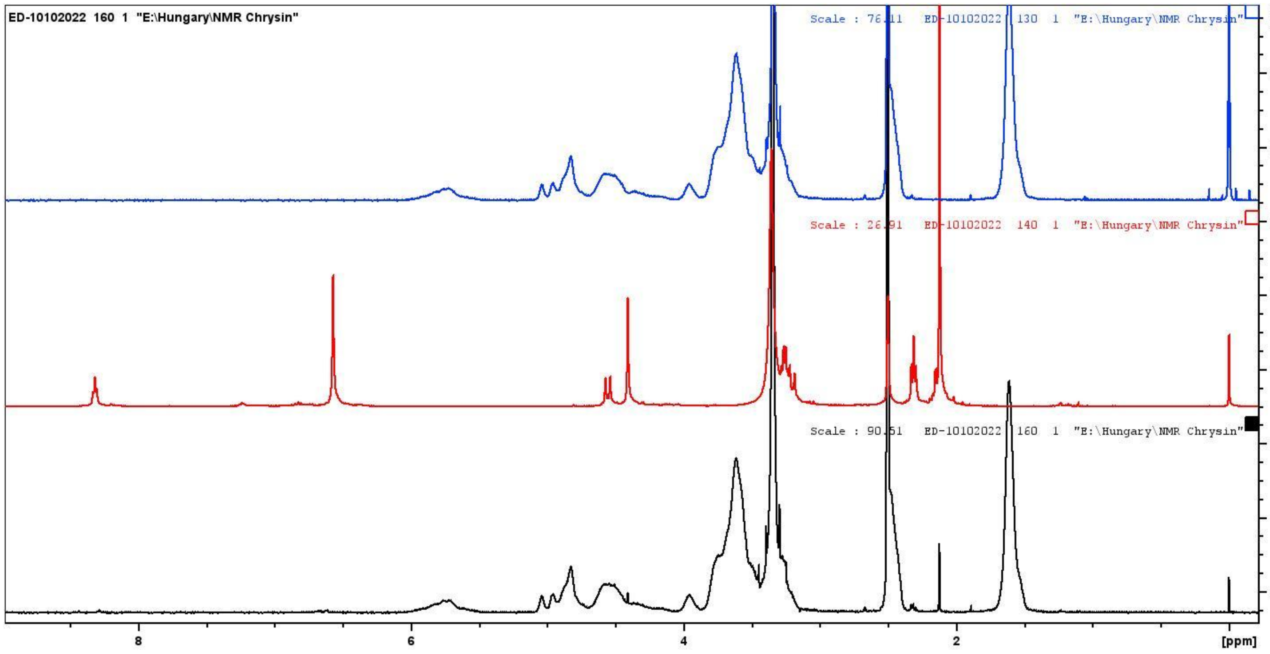


Figure S8. [1]H NMR spectra of OTX008-SBECD mixture (Black line), OTX008 (Red line) and SBECD (Blue line) in DMSO-d6.

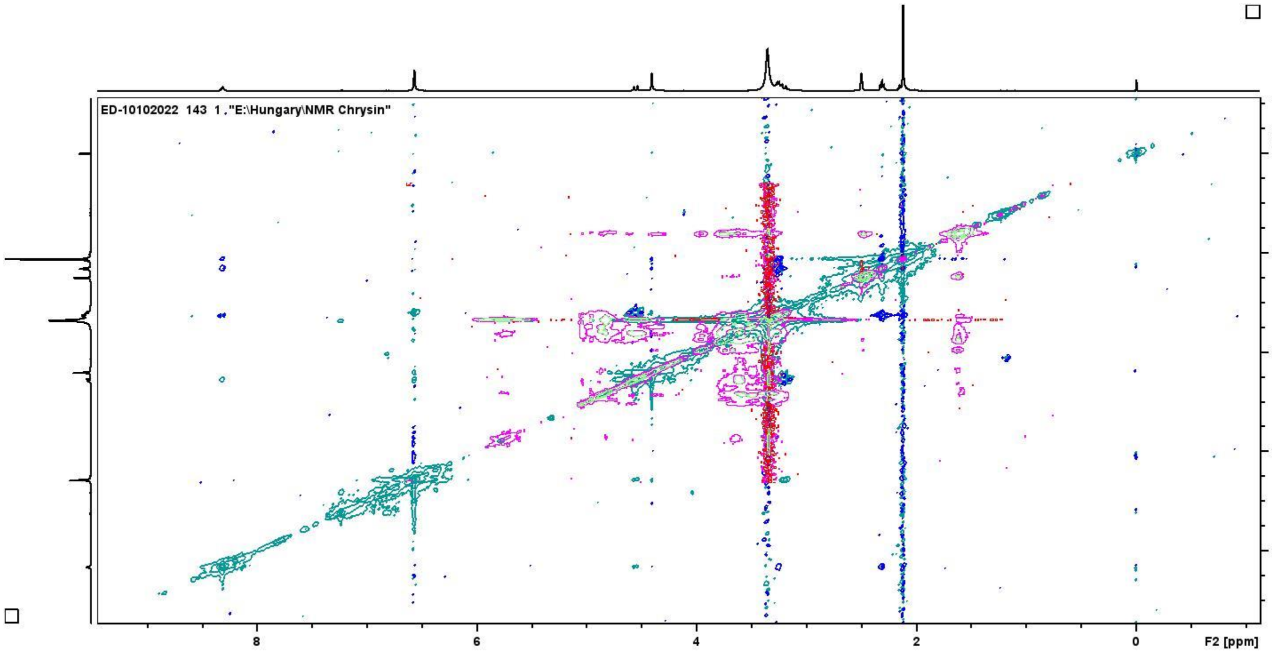


Figure S9. NOESY spectra of OTX008-SBECD mixture (Black line), OTX008 (Red line) and SBECD (Blue line) in DMSO-$d_6$.

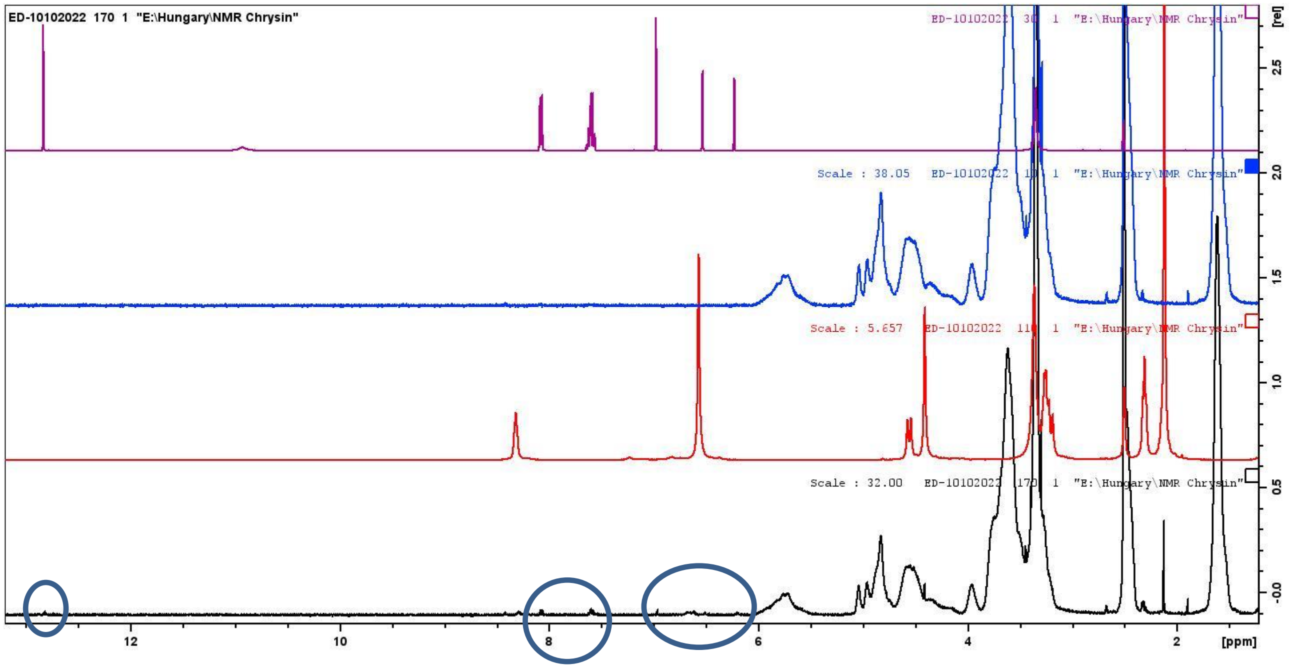


Figure S10. $^1$H NMR spectra of CHR-OTX008-SBECD mixture (Black line), OTX008 (Red line) SBECD (B, Blue line) and CHR (Purple line) in DMSO-$d_6$.

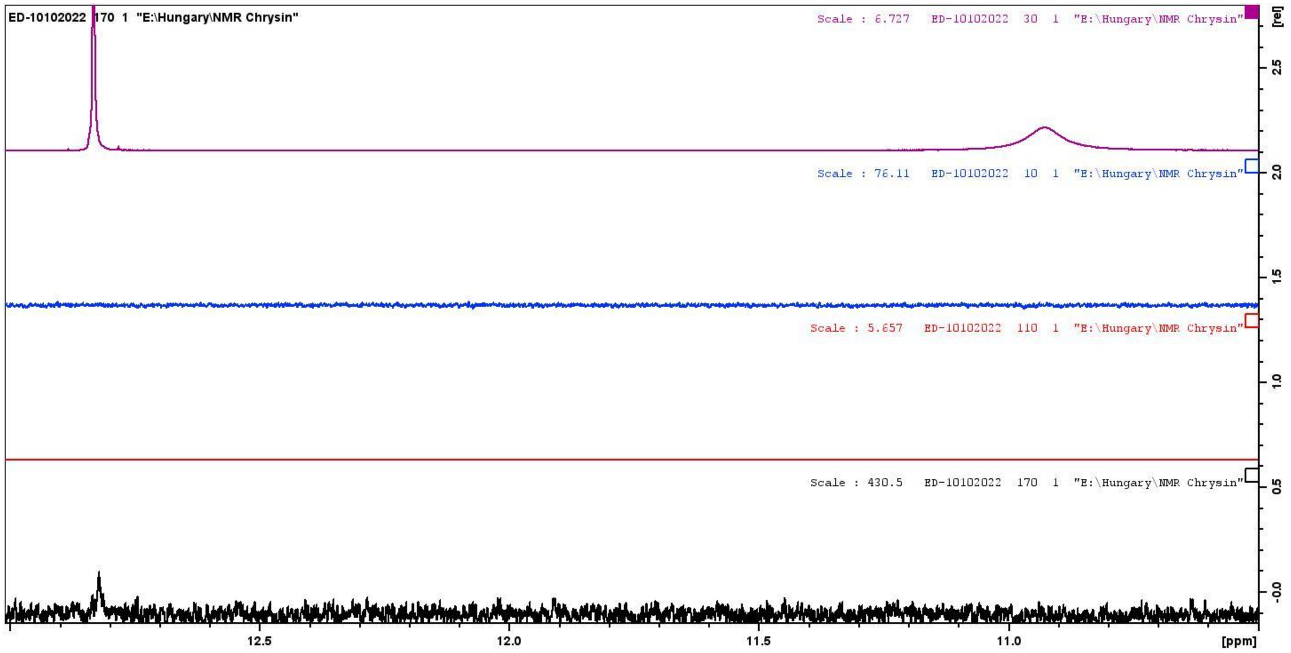


Figure S10a. Expanded area of $^{1}H$ NMR spectra of CHR-OTX008-SBECD mixture (Black line), OTX008 (Red line) SBECD (B, Blue line) and CHR (Purple line) in DMSO-$d_6$.

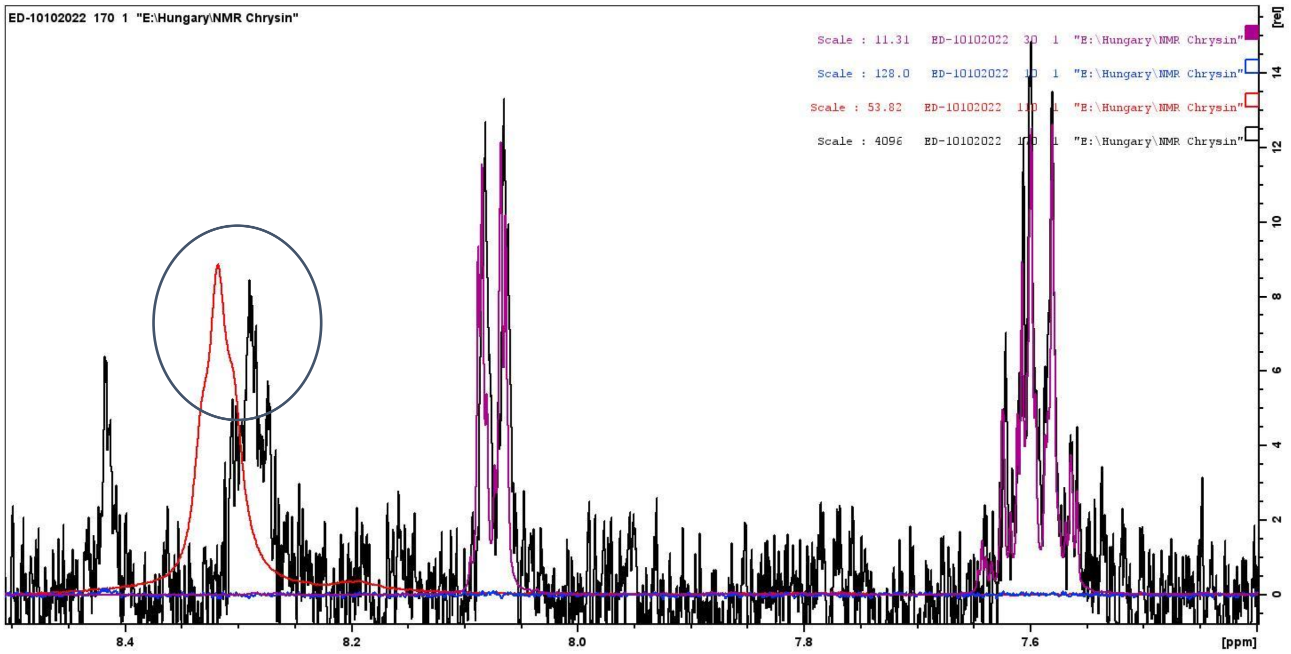


Figure S10b. Expanded area of $^{1}H$ NMR spectra of CHR-OTX008-SBECD mixture (Black line), OTX008 (Red line) SBECD (B, Blue line) and CHR (Purple line) in DMSO-$d_6$.

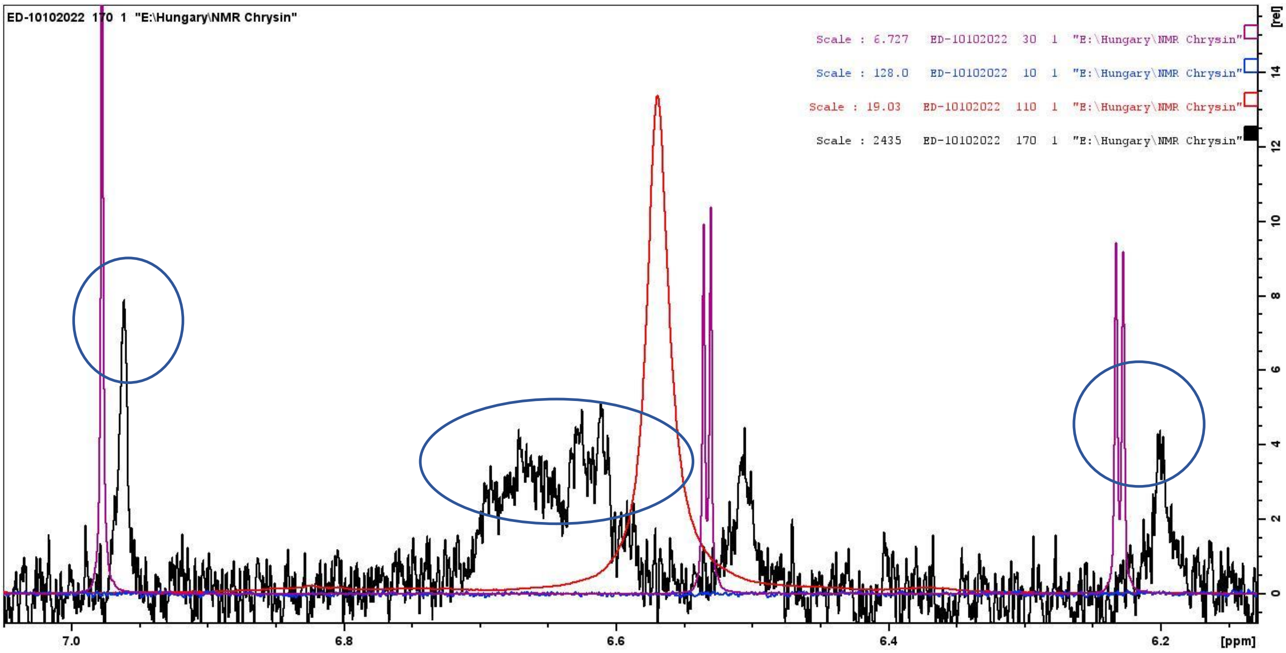


Figure S10c. Expanded area of [1]H NMR spectra of CHR-OTX008-SBECD mixture (Black line), OTX008 (Red line) SBECD (B, Blue line) and CHR (Purple line) in DMSO-$d_6$.

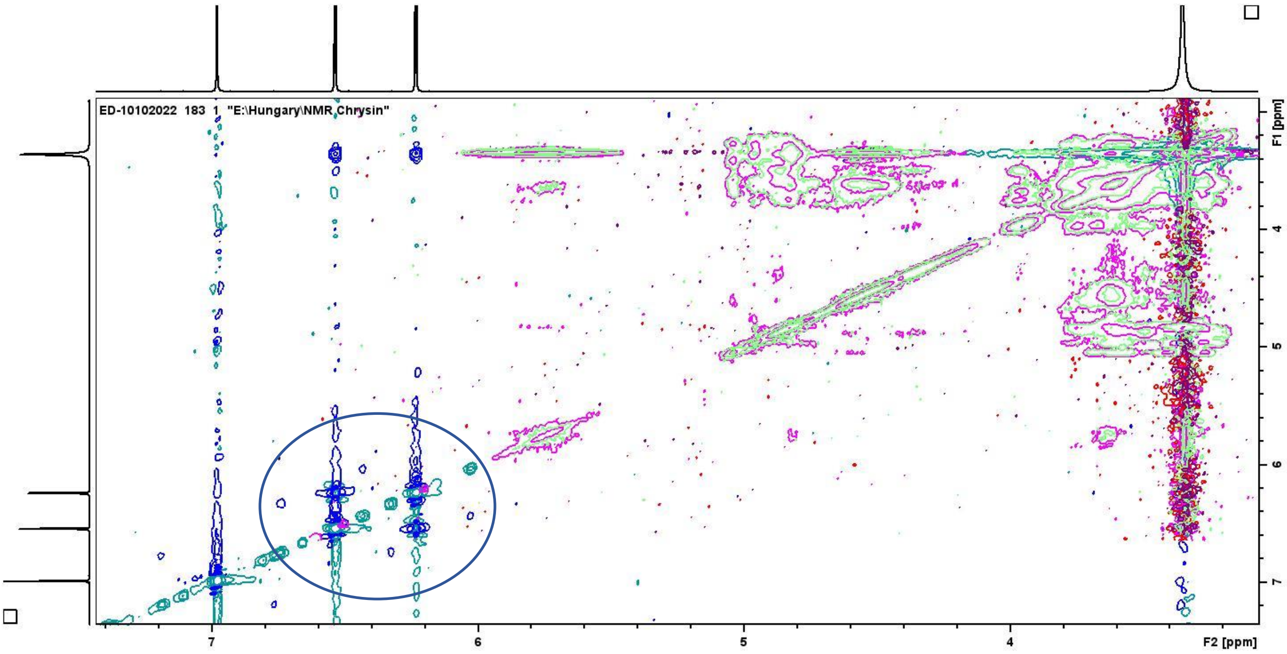


Figure S11. Comparison of NOESY spectra of CHR (Blue/Dark green line), CHR-OTX008-SBECD mixture (purple line) and binary CHR-SBECD mixture (Light green line) in DMSO-$d_6$. The framed parts of the spectrum shown the presense of CHR molecules in a different situation when it is in ternary mixture.

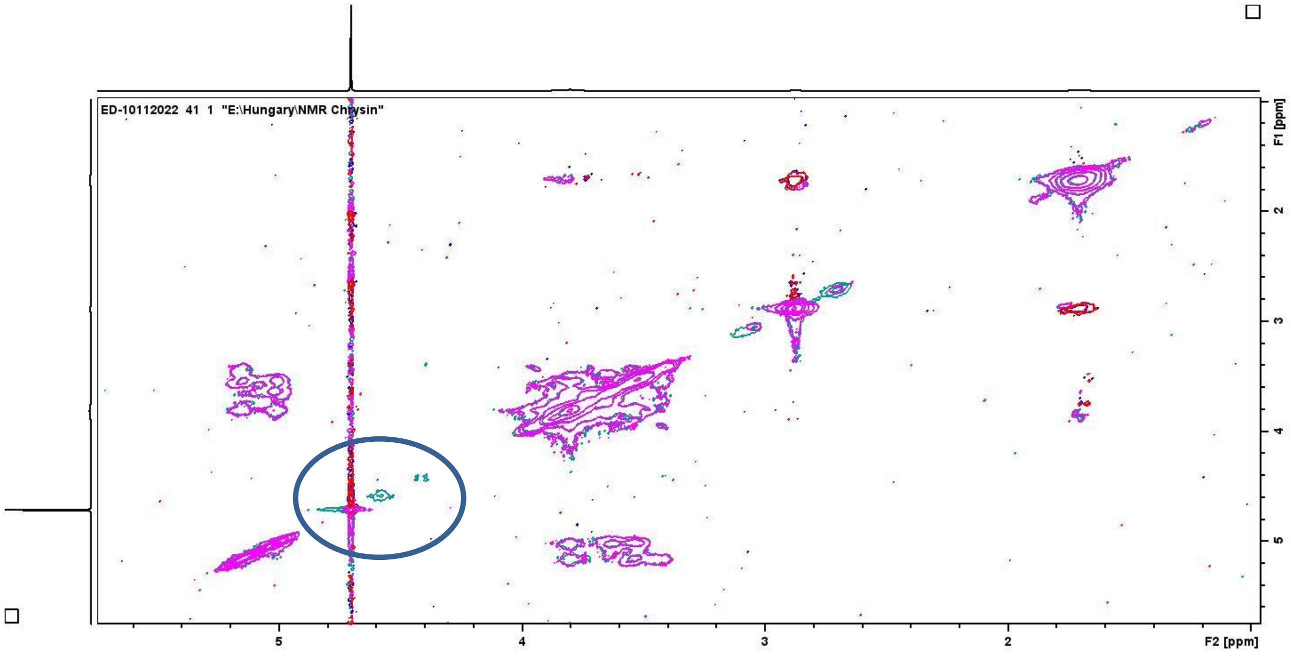


Figure S12. NOESY spectra CHR-OTX008-SBECD mixture (Black line) and SBECD (Red line) in $D_2O$ showing the presense of CHR and OTX008 in the ternary structure.

Table S1. Results of thermal analysis.

| Samples and mixtures | $T_m$ | $T_d$ |
|---|---|---|
| CHR | 290 | - |
| OTX1008 | 218 | 306 |
| SBECD | 272 | 281 |
| Binary CHR- SBECD | 272, 362 | 281 |
| Binary OTX008- SBECD | 290, 362 | 299 |
| Ternary mixture<br>CHR- OTX008- SBECD | 293 | 300 |

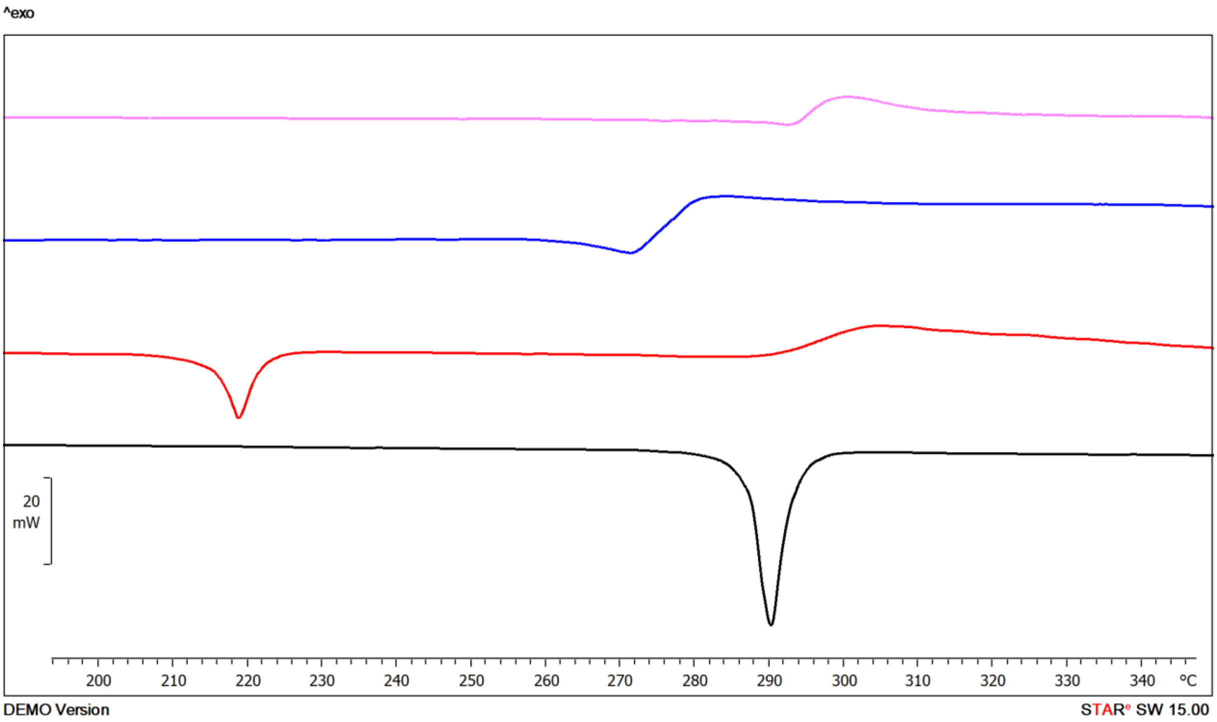


Figure S13. Zoomed view of DSC thermograms of CHR (Black line), OTX008 (Red line), SBECD (Blue line) CHR-OTX008-SBECD (Purple line), at 190-350 °C, 10 °C/min, inert $N_2$.

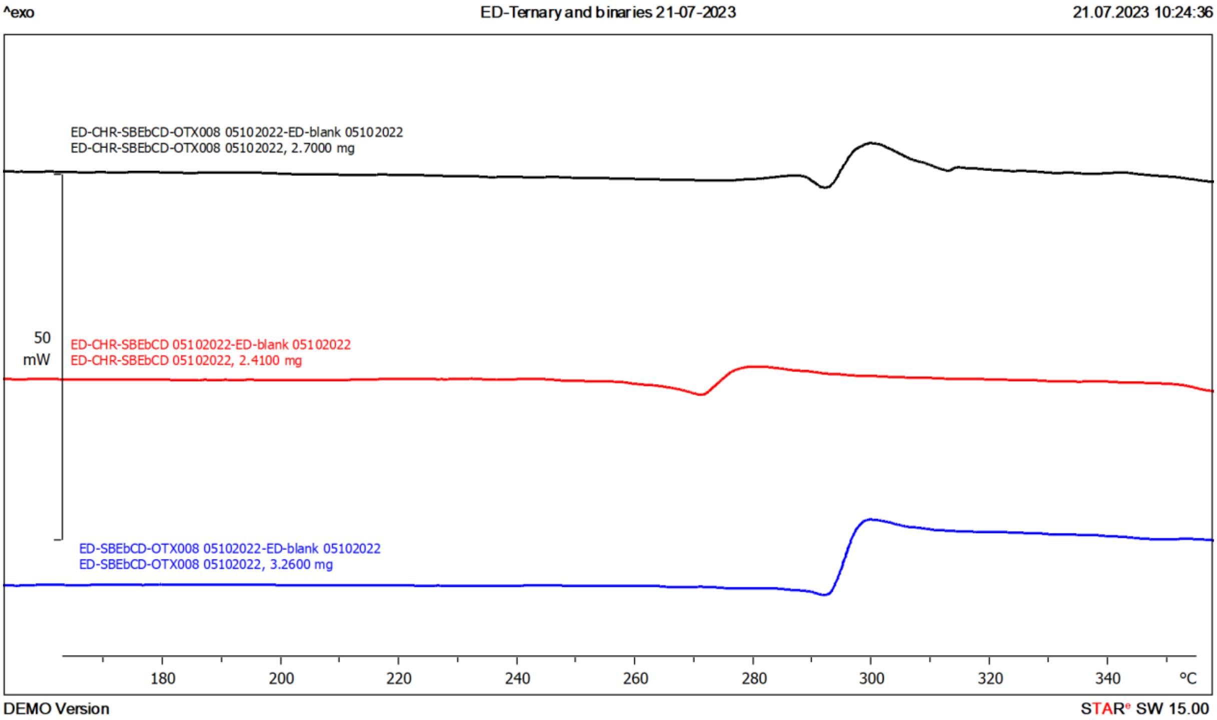


Figure S14. Comparison of zoomed areas of DSC thermograms of SBECD-OTX008 mixture (Blue line), SBECD-CHR mixture (Red line) and CHR-SBECD-OTX008 mixture (Black line), at 160-350 °C, 10 °C/min, inert $N_2$.

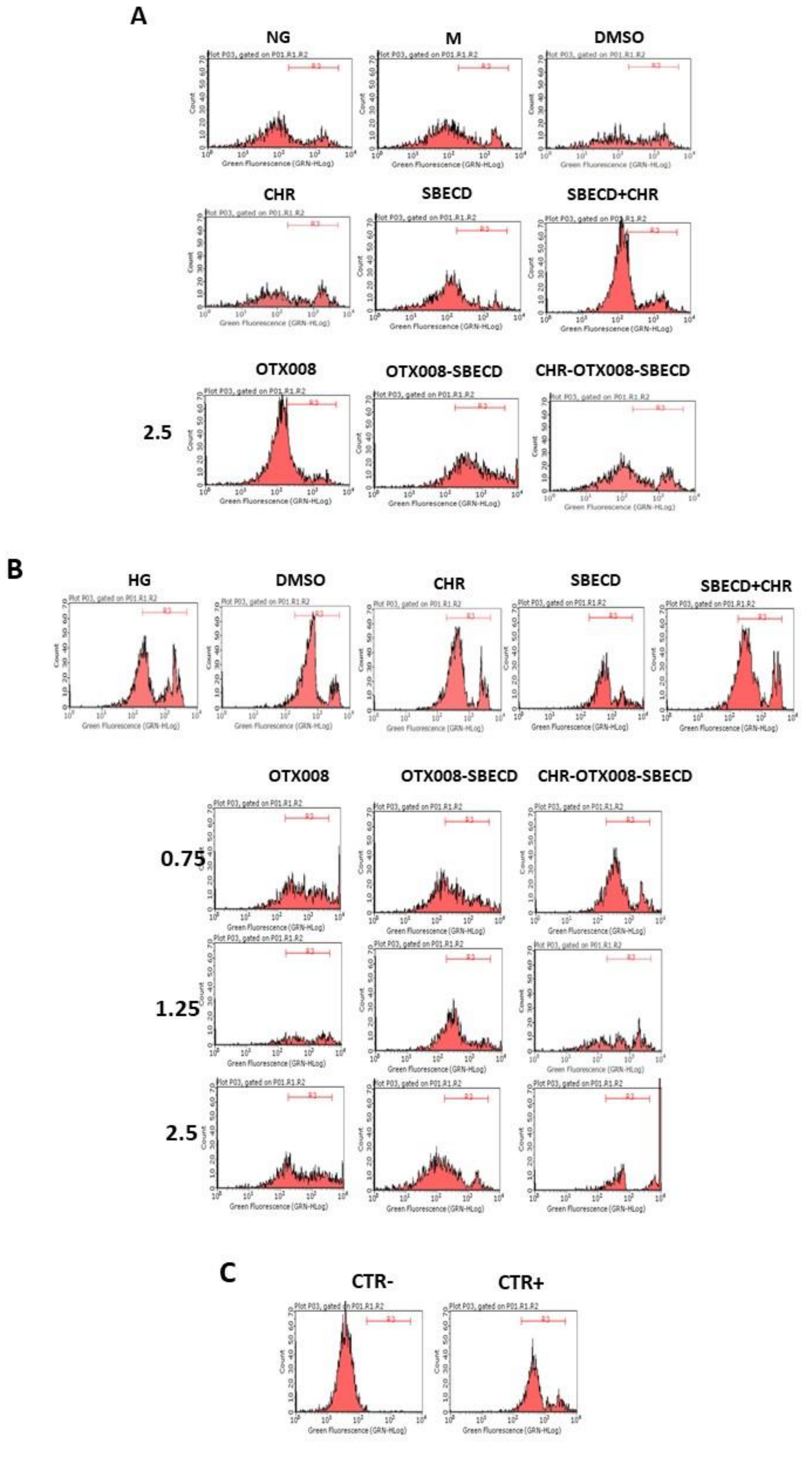


Figure S15. (A) Representative flow cytometer measures of total intracellular ROS levels assayed with DCFH-DA probe in NG or (B) HG medium. In NG cells, OTX008 was tested at the maximum dose of 2.5 μM. In HG cells, OTX008 was tested at the doses of 2.5-1.25-0.75 μM; (C) CTR– = negative control (5% FBS without DCFH-DA); CTR+= positive control (H2O2 100 μM). R3 region = DCFH-DA-positive cells.

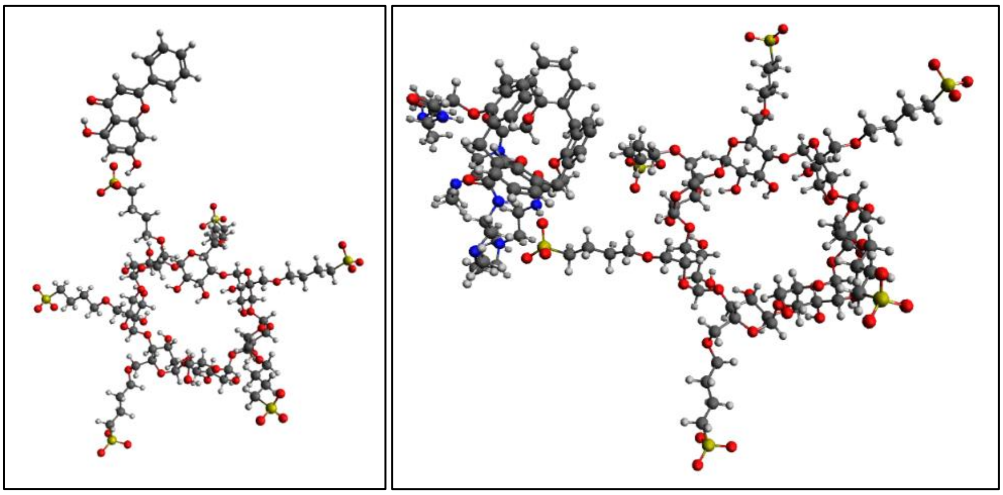

Figure S16. Interactions between SBECD and CHR (A). Interaction between OTX008 and SBECD (B).
